# Broad-spectrum polerovirus resistance conferred by a potato TIR-NLR immune receptor

**DOI:** 10.64898/2026.06.29.735250

**Authors:** Robert Heal, He Zhao, Hee-Kyung Ahn, Maria Sindalovskaya, John Walsh, Jan Kreuze, Hannele Lindqvist-Kreuze, Kamil Witek, Jonathan D G Jones

**Affiliations:** The Sainsbury Laboratory, University of East Anglia, Norwich Research Park, Colney Lane, Norwich, NR4 7UH, UK; Institute of Molecular Plant Sciences, University of Edinburgh, Daniel Rutherford Building, King’s Buildings, EH9 3BF, Edinburgh, UK; John Innes Centre, Norwich Research Park, Colney Lane, Norwich, NR4 7UH, UK; School of Life Sciences, University of Warwick, Wellesbourne Campus, Warwick, CV35 9EF, UK; Genetics Genomics and Crop Improvement, International Potato Center, Lima, Peru; 62Blades, Evanston, IL, 60201, USA

## Abstract

Potato leafroll virus (PLRV) is an economically important viral disease of potato (*S. tuberosum*). Genetic resistance to this phloem-limited virus is rare, and no cloned resistance (*R*) genes have been reported. *Rl_adg_* confers resistance to PLRV in an Andean potato landrace, LOP-868 (Velásquez et al. 2007). We identified the functional *Rl_adg_* gene as a homolog of the tomato TIR-NLR-encoding *Bs4*. Rl*_adg_* interacts with the serine protease domain of the PLRV protein P1, which is essential for virus replication. This recognition is independent of the protease’s enzymatic activity, and the Rl_adg_ immune receptor oligomerizes upon direct association with the protease. Like PLRV, many poleroviruses contain a serine protease. Despite their diverse amino acid sequences, these proteases are predicted to share similar structures. Rl_adg_ recognizes all ten tested polerovirus proteases, suggesting a conserved structural recognition mechanism. We propose that Rl_adg_’s broad recognition capacity could enable resistance to poleroviruses in many crop species. *Rl_adg_* is the first *R*-gene reported to confer resistance to a phloem-limited pathogen and could provide enhanced resistance to many economically important poleroviruses.

## Introduction

Plant viruses form a complex polyphyletic group that interact not only with their primary plant hosts but also with additional hosts and vectors, including nematodes, arthropods, oomycetes, fungi, and protists (Lefeuvre et al., 2019, Tamada and Kondo, 2013, Andika et al., 2017, Andret-Link et al., 2004). They can exhibit a range of host specificity, from infecting a single plant species to having a broad host range, and their effects can vary from highly symptomatic infections to those that appear symptomless.

*Polerovirus* is a genus of plant-infecting viruses within the *Solemoviridae* (formerly *Luteoviridae*) family. Poleroviruses are phloem-limited and are transmitted via aphid vectors in a circulative, persistent manner. While they replicate in the plant host phloem, they do not in the insect vector (Macleod et al., 2023, Ryabov et al., 2026). This genus includes economically important viruses such as turnip yellows virus (TuYV), beet mild yellowing virus (BMYV) and pepper vein yellows virus (PeVYV), which infect oilseed rape, sugar beet, and pepper, respectively. The Polerovirus type species is potato leafroll virus (PLRV), a viral pathogen of potato. PLRV infection can cause yield losses of 50-80% and reduce tuber quality through the development of net necrosis (Kondrák et al., 2020, Carneiro et al., 2017). While primary infection may produce only mild symptoms, shoots grown from infected tubers can become stunted and discoloured. Co-infection with other viruses often exacerbates symptoms, potentially due to the loss of phloem restriction observed in co-infections with potato virus A and Y (Hameed et al., 2014, Jingwei et al., 2013, Barker, 1987, Savenkov and Valkonen, 2001).

Poleroviruses carry a positive-sense single-stranded RNA (ssRNA) genome with a conserved architecture with five major open reading frames (ORFs: ORF0-ORF5). ORF0 encodes P0, a suppressor of RNA silencing. The ORF1-2 region encodes several replicase-associated proteins; Rap1, the P1 protein and a P1-P2 fusion protein formed via a -1 ribosomal frameshift from the P1 ORF. ORF4 encodes a movement protein while ORF3 and ORF5 encode two components of the coat protein (CP), with ORF5 being produced through a readthrough of the ORF3 stop codon (Nurkiyanova et al., 2000, Taliansky et al., 2003).

Currently, PLRV is managed by targeting its insect vectors, mostly *Myzus persicae*, using broad-spectrum insecticides. However, the growing prevalence of insecticide resistance, coupled with the effects of climate change and withdrawal of approval for insecticide use, is already intensifying the impact of *M. persicae*-vectored viruses especially PLRV (Ryabov et al., 2026). Warmer winters may lead to earlier aphid flights in spring and later flights in autumn and allow the vector population to build up on alternative hosts before the crop emergence (Hemming et al., 2022). Enhancing crop immunity offers a sustainable, chemical-free solution to combat plant viruses.

Intracellular pathogen recognition in plants is primarily mediated by the nucleotide-binding leucine-rich repeat (NLR) class of immune receptor proteins, often encoded by *Resistance* (*R-*) genes. Plant NLRs are usually classified into three groups based on their N-terminal domain, ‘Toll, interleukin-1 receptor’ (TIR-NLR), ‘Coiled-coil’ (CC-NLR) or ‘Resistance to powdery mildew 8’ (RPW8-NLR). All three groups share a central NB-ARC domain and a C-terminal LRR domain (Feehan et al., 2020). Recent phylogenomic analyses reveal high diversity within the CC-NLR clade (Contreras et al., 2023).

Several *R*-genes conferring resistance to plant viruses have been cloned, many of which encode NLR proteins such those for resistance to potato virus X (*Rx-1*, *Rx-2*) and potato virus Y (*Ry_sto_*) (Bendahmane et al., 1999, Bendahmane et al., 2000, Grech-Baran et al., 2019). However, no NLR-encoding *R*-genes conferring resistance to PLRV or other poleroviruses have been reported. It remains unclear whether conventional NLR-mediated effector-triggered immunity functions in the phloem (Bendix and Lewis, 2018, Jiang et al., 2019). Notably, NLR-encoding transcripts have been detected in phloem companion cells and procambium (Tang et al., 2023). Since poleroviruses move between sieve elements, companion cells and vascular parenchyma cells (Esau and Hoefert, 1972, Shepardson et al., 1980), NLR-mediated immunity could play a role in responding to these pathogens. Genes conferring resistance to turnip yellows virus (TuYV) have been identified in *Brassica rapa*, *Brassica oleracea* and *Brassica napus*; these genes (which have not been isolated) are only partially dominant and do not exhibit major effects (Greer et al., 2021, Hackenberg et al., 2020, Juergens et al., 2010, Macleod et al., 2023). Resistance to both TuYV and PLRV has been reported in *Nicotiana glutinosa*, where this trait is monogenic and segregates in a 3:1 ratio in biparental crosses. Expression of the PLRV P0 protein elicits a response in the resistant *N. glutinosa* accession. This *R*-gene, named *RPO1* (*Resistance to poleroviruses 1*), has yet to be cloned (Wang et al., 2015, Wang et al., 2023).

Through extensive screening, Mihovilovich et al. (2007) identified three *S. tuberosum ssp. andigena* landraces with high levels of heritable resistance to PLRV. These accessions - LOP-868, HUA-332 and OCH-7643 – demonstrate strong resistance, with viral titres remaining consistently low following graft inoculation with PLRV-infected plants. Using a dihaploid mapping population, Velasquez et al. (2007) mapped resistance in LOP-868 to a single major locus, *Rl_adg_,* on the long arm of Chromosome 5. This region contains numerous NLR-encoding genes (Jupe et al., 2012, Kuang et al., 2005). The use of *Rl_adg_* in breeding programmes is currently limited by the complexities of tetraploid potato genetics and the absence of a perfect molecular marker. Furthermore, crosses between LOP-868 and other potato cultivars result in progeny that display significant variation in tuber yield, size and appearance (Carneiro et al., 2017). We report here that *Rl_adg_* encodes a TIR-NLR ortholog of *Bs4*, a *Solanum lycopersicum* gene that confers resistance to bacterial spot disease. Rl_adg_ directly recognises a serine protease domain found in PLRV and other poleroviruses.

## Results

### Transient expression of the PLRV P1-P2 genomic region elicits HR in potato containing *Rl_adg_*

We set out to isolate *Rl_adg_*, the gene responsible for PLRV resistance in the landrace potato variety LOP-868. Testing *S. tuberosum* for resistance to PLRV is challenging, as it requires either grafting infected tissue or the transfer of aphids carrying the virus. To simplify candidate gene evaluation, we identified the PLRV protein recognised by Rl_adg_. To achieve this, we amplified ORFs from a PLRV cDNA clone and cloned them into a CaMV *35S* expression vector. The constructs were designed to transiently express either P0, the P1-P2 region (encompassing Rap1, P1 and the P1-P2 fusion protein), the movement protein (MP), or the coat protein (CP) (Fig. 1a). These constructs were transiently expressed in the leaves of LOP-868 (the resistant accession) and the PLRV-susceptible cultivar Maris Piper to identify the eliciting protein. Expression of the P1-P2 construct in LOP-868 resulted in strong HR, indicating that the elicitor of Rl_adg_ is either P1, the P1-P2 fusion protein, or Rap1. In contrast, no response was observed for any other PLRV ORF in LOP-868 or Maris Piper (Fig. 1b). This approach enabled the development of a rapid assay for identifying functional *Rl_adg_* paralog by co-expression with the P1-P2 domain.

**Figure 1.**
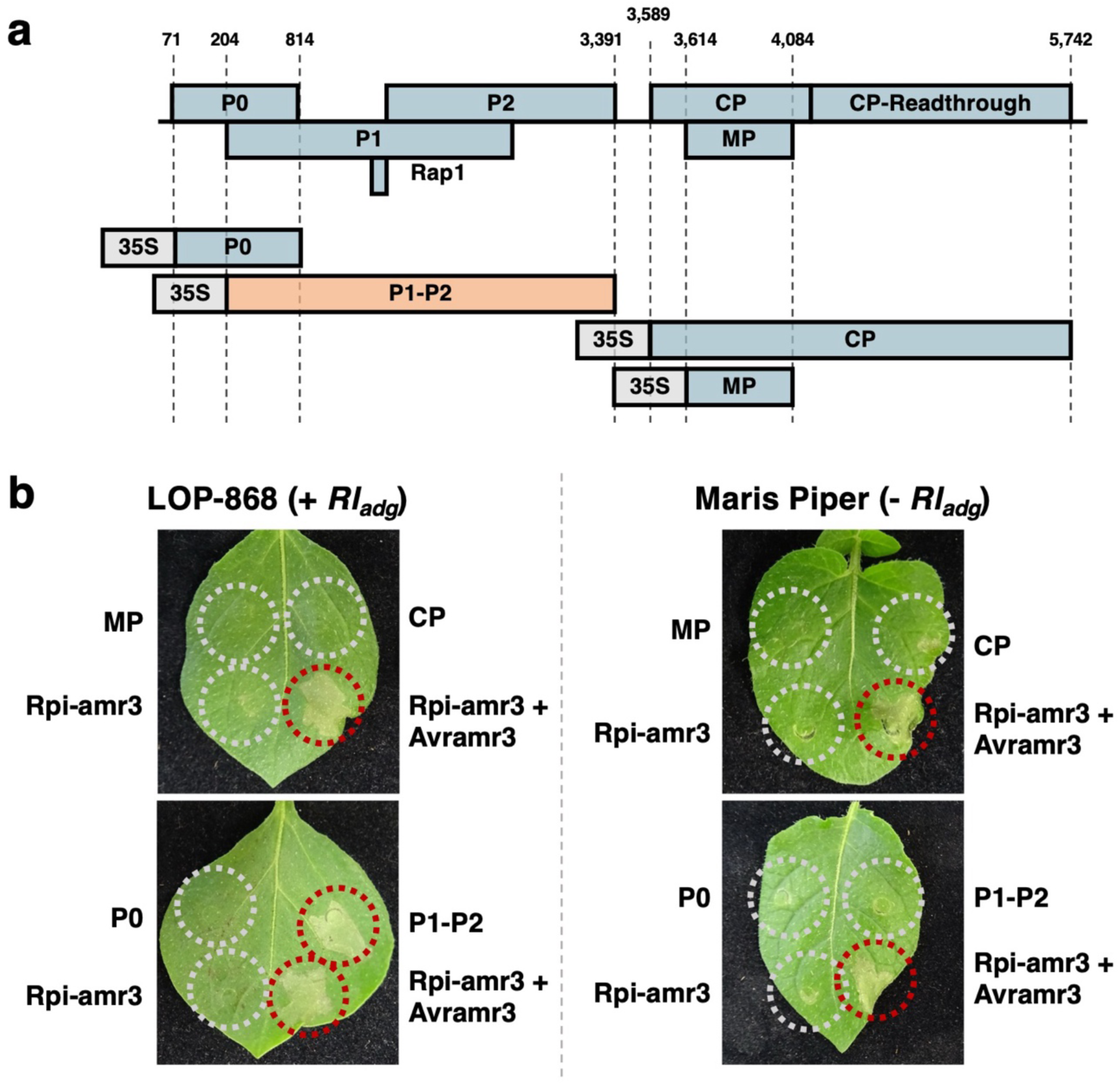
Transient expression of the PLRV P1-P2 genomic region elicits HR in potato containing *Rl_adg_*. **(a)** To determine which PLRV protein is recognized by Rl_adg_, the viral genome was divided into four constructs (P0, P1-P2, CP, and MP), each placed under the control of the CaMV 35S promoter and Ocs terminator. Construct boundaries within the PLRV genome (GenBank:KP090166.1) are indicated. **(b)** Constructs were transiently expressed in leaves of LOP-868 (which carries *Rl_adg_*) and the potato cultivar Maris piper (which lacks *Rl_adg_*) by *Agrobacterium* infiltration. Expression of the P1-P2 region induces an *Rl_adg_*-dependent HR. *Agrobacterium* (Agl1 strain) suspensions were adjusted to OD_600_ = 0.1 prior to infiltration. Photographs were taken at 3 days post-infiltration.

### Two classes of NLR-encoding gene co-segregate with *Rl_adg_* in a dihaploid population

To identify *Rl_adg_*, we utilised PacBio RenSeq to assemble the NLR repertoire of LOP-868, and conducted bulked segregant analysis on a phenotyped dihaploid population to identify alleles present in the resistant haplotype. Illumina RenSeq was performed on bulked resistant and susceptible plants, each containing DNA from 30 plants from the dihaploid population phenotyped by Velasquez et al. (2007). Reads from the bulked segregant datasets were mapped to the LOP-868 SMRT RenSeq assembly and resistance-linked sequences were identified by searching for contigs where read depth was reduced in the susceptible dataset but maintained in the resistant dataset. In LOP-868, 27 *R1* orthologs and 17 *Bs4* orthologs were identified. Of these, eight *R1* orthologs and three *Bs4* orthologs were linked to the resistant haplotype (Fig. 2a) (Supplementary Fig. 1). To further refine the candidate list, cDNA RenSeq was performed on leaf tissue from LOP-868 to determine gene expression. This analysis confirmed the expression of six *R1* orthologs and three *Bs4* orthologs (Supplementary Table.1) (Fig. 2a). These nine genes were selected for further testing.

**Figure 2.**
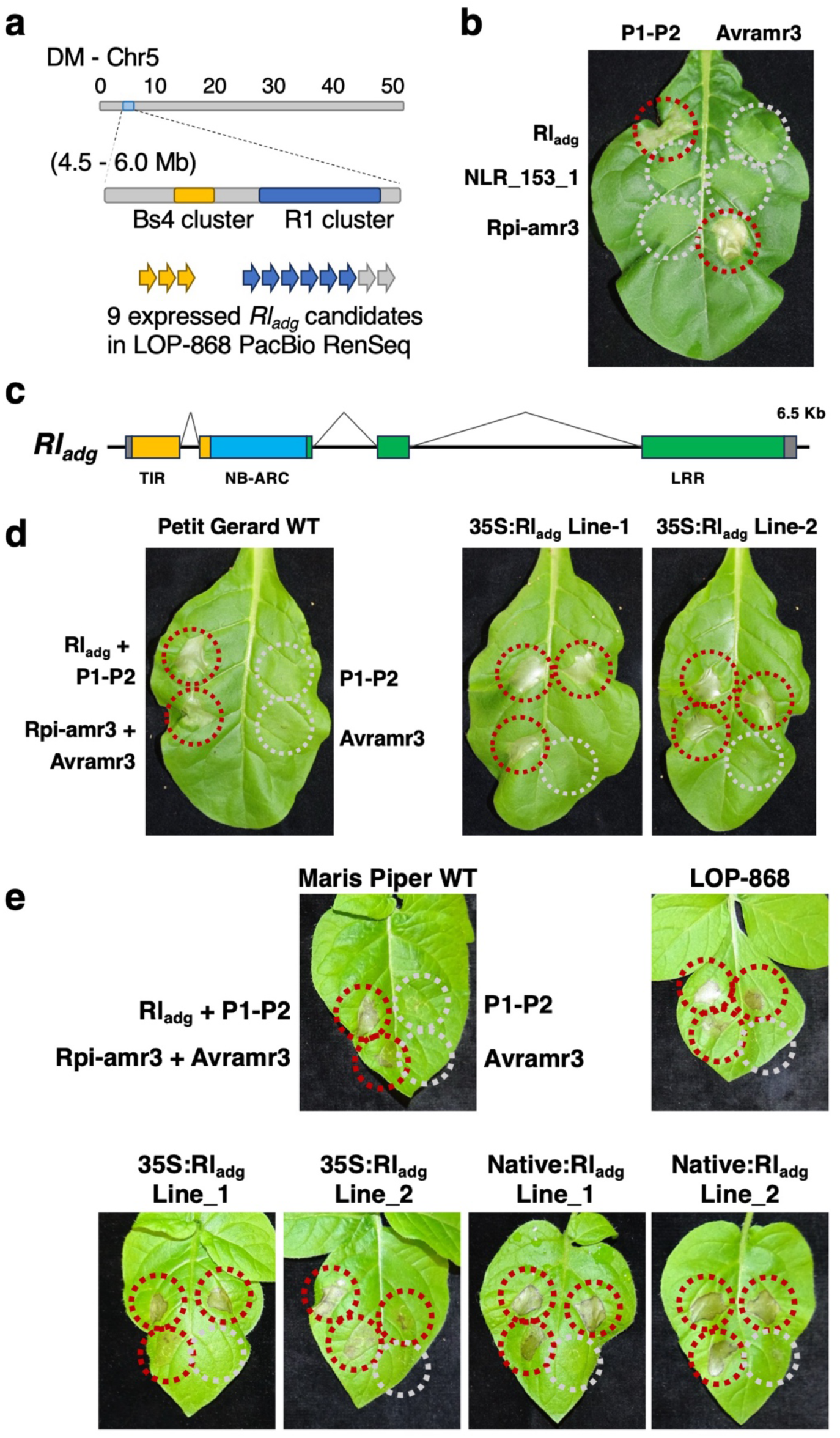
*Rl_adg_* is a *Bs4* orthologue located on *S. tuberosum* Chromosome 5. **(a)** In a previously characterized dihaploid population (LOP-868 x IVP-101; Velasquez et al. 2007), *Rl_adg_* was mapped to the Chromosome 5 *R1* and *Bs4 NLR* gene clusters. DNA from 30 resistant and 30 susceptible dihaploids was pooled into separate RenSeq libraries. Bulked segregant analysis identified *R1* and *Bs4* orthologs specific to the resistant haplotype. cDNA RenSeq revealed three expressed *Bs4* orthologs and six expressed *R1* orthologs as candidates. **(b)** Two of the three resistance-linked *Bs4* homologs were cloned from LOP-868. Transient expression assays in *N. tabacum* showed that co-expression of *Rl_adg_* with *P1-P2* triggered HR. In contrast, no cell death was observed when *Rl_adg_* was co-expressed with *Avramr3*, or when *NLR_153_1* (a non-functional *Rl_adg_* paralog) or *Rpi-amr3* were expressed with *P1-P2*. *Agrobacterium* (Agl1 strain) was infiltrated at OD_600_ = 0.3. Photographs were taken 3 days post-infiltration. **(c)** *Rl_adg_* encodes a 1,123-amino acid TIR-NLR sharing 79.8% identity with Bs4. Domains are indicated as follows: TIR (yellow), NB-ARC (blue), and LRRs (green). The 5’ and 3’ UTRs are shown in grey. Intron-exon boundaries were defined using cDNA RenSeq data, with no evidence of alternative splicing. **(d)** Transient expression of P1-P2 in *N. tabacum* lines expressing *Rl_adg_* induced HR. *Agrobacterium* (Agl1 strain) was infiltrated at OD_600_ = 0.3. Photographs were taken at 3 days post-infiltration. **(e)** Transgenic *S. tuberosum* (cv. Maris Piper) expressing *Rl_adg_* also recognize P1-P2, triggering HR. *Agrobacterium* (Agl1 strain) was infiltrated at OD_600_ = 0.1, and photographs were taken at 3 days post-infiltration.

### A Bs4-homologous TIR-NLR confers responsiveness to PLRV P1-P2

Of the nine NLRs identified as *Rl_adg_* candidates, we successfully cloned six *R1* orthologs and two *Bs4* orthologs. The candidates were co-expressed with the PLRV P1-P2 region in *Nicotiana tabacum*. No cell death was observed with any *R1* homolog (Supplementary Fig. 2). However, co-expression of *P1-P2* with one *Bs4* ortholog resulted in a strong hypersensitive response (Fig. 2b). This gene (*NLR_97_1*) is *Rl_adg_*. No cell death was observed when the second cloned *Bs4* ortholog (*NLR_153_1*) was expressed with *P1-P2*. *Rl_adg_* is a four-exon gene that encodes a 1,123 amino acid TIR-NLR (Fig. 2c), that shares 79.8% amino acid identity with Bs4.

To confirm that *Rl_adg_* alone is sufficient for recognition of PLRV, we generated transgenic *S. tuberosum* and *N. tabacum* lines. Constructs containing the *Rl_adg_* ORF, driven by either its native promoter or the CaMV *35S* promoter, were transformed into *S. tuberosum*. In parallel, transgenic *N. tabacum* lines expressing *Rl_adg_* under the CaMV 35S promoter were also produced. Expression of the PLRV P1-P2 region in the transgenic *N. tabacum* lines was sufficient to trigger HR (Fig. 2d). Similarly, both native and CaMV *35S*-driven constructs conferred recognition of P1-P2 in transgenic potato plants (Fig. 2e, Supplementary Fig. 3). These findings demonstrate that the delivery of a single NLR gene, *Rl_adg_*, is sufficient to reconstitute the LOP-868 HR phenotype in *S. tuberosum* and *N. tabacum*.

### Rl_adg_ recognises the serine protease domain of P1, independent of protease function

The PLRV P1-P2 genomic region encodes three proteins: P1, Rap1, and the P1-P2 fusion protein (Fig. 3a). To determine which of these proteins is recognised by Rl_adg_, we truncated the P1-P2 construct, removing either the P1 or P2 ORF. Additionally, the Rap1 start codon was mutated. Transient expression assays in *N. tabacum* confirmed that P1 alone is sufficient to trigger HR when co-expressed with *Rl_adg_* (Fig. 3b).

**Figure 3.**
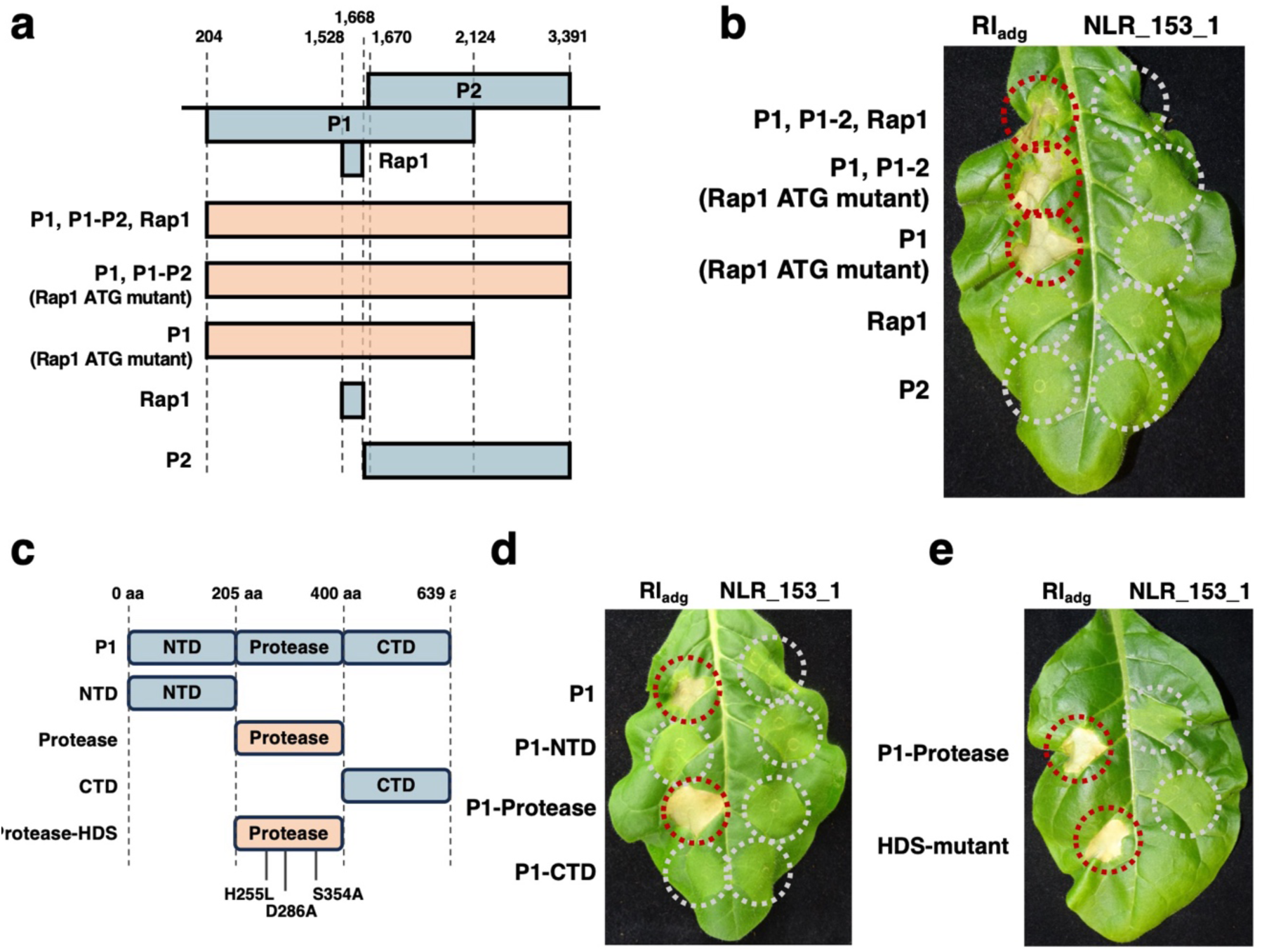
Rl_adg_ recognizes the protease domain of P1, independent of its activity. **(a)** The PLRV P1-P2 genomic region encodes three proteins: P1, a P1-P2 fusion protein, and Rap1. To determine which is recognized by Rl_adg_, constructs were generated by truncation or start codon mutation. The relative positions of the ORFs within the PLRV reference genome (GenBank:KP090166.1) are shown. Constructs that trigger an Rl_adg_-dependent response in *N. tabacum* are highlighted in red. **(b)** In *N. tabacum,* expression assays showed that neither Rap1 nor P2 are required for Rl_adg_-dependent cell death; the P1 ORF alone is sufficient. The non-functional *Rl_adg_* paralog *NLR_153_1* was used as a negative control. *Agrobacterium* (Agl1 strain) was infiltrated at OD_600_ = 0.3. Images were taken 3 days post-infiltration. **(c)** P1 undergoes self-cleavage via serine protease activity, producing three fragments: an N-terminal domain, a serine protease, and a C-terminal domain. Truncated constructs expressing each fragment individually were generated, along with a mutant protease containing substitutions in the HDS catalytic triad (amino acid changes were replicated from Sadowy et al., 2001). **(d)** The protease domain alone is sufficient for Rl_adg_ activation, while no cell death was observed following expression of the N-terminal or C-terminal fragments. The non-functional *Rl_adg_* paralog *NLR_153_1* was used as a negative control. *Agrobacterium* (Agl1 strain) was infiltrated at OD_600_ = 0.3. Photographs were taken 3 days post-infiltration. **(e)** The HDS catalytic triad mutant of the P1 protease elicited cell death when co-expressed with *Rl_adg_*, at levels equivalent to the functional protease. The non-functional *Rl_adg_* paralog *NLR_153_1* was used as a negative control. *Agrobacterium* (Agl1 strain) was infiltrated at OD_600_ = 0.3. Photographs were taken 3 days post-infiltration.

P1 is a polyprotein that self-cleaves into three proteins: an N-terminal domain (NTD) of unknown function, a central serine protease domain, and a C-terminal domain (CTD) which acts as a viral genome-linked protein (Vpg) capping the RNA (Fig. 3c). To pinpoint the region recognised by Rl_adg_, we further truncated P1 into its constituent domains, each expressed under the CaMV 35S promoter. Co-expression of *Rl_adg_* with the serine protease domain elicited HR in *N. tabacum*, whereas neither the NTD nor the CTD triggered a response (Fig. 3d), thus revealing the serine protease domain as the elicitor of Rl_adg_-mediated recognition.

To evaluate whether this recognition depends on protease activity, we engineered a non-functional variant of the serine protease. The catalytic activity of serine proteases relies on an HDS triad (His254, Asp286, Ser353), and the mutation of any of these residues abolishes functionality (Sadowy et al., 2001). We generated an HDS-mutant (H254L-D286A-S353A), which, when co-expressed with *Rl_adg_*, still triggered strong HR comparable to the functional protease (Fig. 3e). These results demonstrate Rl_adg_ recognises the serine protease domain independently of its catalytic function. This suggests that recognition occurs via direct interaction with the protease, rather than detecting its activity or a processed host target.

### *Rl_adg_* confers recognition of a diverse set of poleroviruses

Many NLRs recognise conserved elicitor structures shared across pathogens, and this recognition can translate into broad-spectrum resistance. For example, while the *S. stoloniferum* TIR-NLR Ry_sto_, originally identified as a potato virus Y (PVY) resistance protein, also recognises the coat proteins of several other potyviruses (Grech-Baran et al., 2019, Grech-Baran et al., 2022).

We identified orthologs of PLRV P1 protease in many solemoviruses, although many show low (40.6% - 82.6%) sequence identity with the PLRV protease (Supplementary Fig. 4). To investigate whether these proteases share similar structures, we used Alphafold to predict ten polerovirus protease structures - including PLRV. The models generated were of high quality, with LDDT (local distance difference test) scores exceeding 80 for most positions within the models (Supplementary Fig. 5). As no structural data exists for any polerovirus serine protease, we evaluated these models by comparing them with the known structure of the sesbania mosaic virus (SeMV) serine protease (Gayathri et al., 2006). SeMV, a member of the *Sobemovirus* genus, shares classification within the *Solemoviridae*. The predicted structures of polerovirus proteases were highly similar to each other, as well as to the SeMV protease (Supplementary Fig. 5). To quantify the structural similarity, we used DALI (Holm et al., 2023). Pairwise z-scores between all polerovirus proteases were high (above 20), indicating strong structural similarity (Supplementary Fig. 6).

Next, we tested whether *Rl_adg_* recognises proteases from other poleroviruses. We synthesised the proteases of nine different poleroviruses and tested them by co-expression with *Rl_adg_* in *N. tabacum*. We observed strong *Rl_adg_*-dependent HR for all nine proteases (Fig. 4a). To determine if this recognition acts against viral infections, we co-expressed *Rl_adg_* and an infectious TuYV cDNA clone in *N. tabacum* leaves. At 4 days post-infiltration, we observed clear cell death (Fig. 4b). Similarly, strong cell death was evident in leaves infiltrated with *Agrobacterium* strains carrying an infectious clone of PLRV (Fig. 4b). These findings suggest that *Rl_adg_* may be capable of elevating resistance to TuYV, and other poleroviruses.

**Figure 4.**
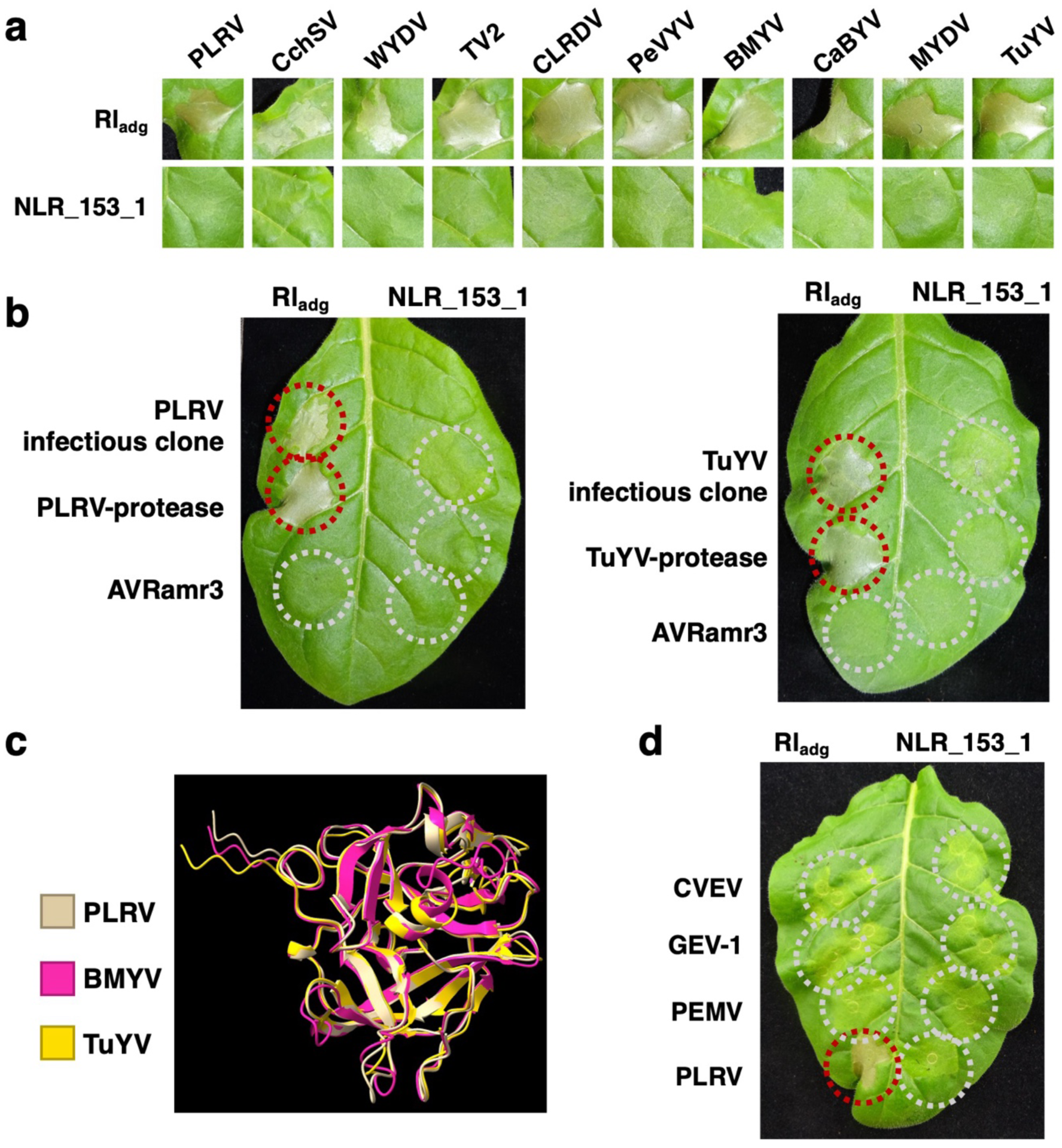
*Rl_adg_* confers recognition of a diverse set of poleroviruses. **(a)** In *N. tabacum* transient assays, Rl_adg_ recognizes at least ten polerovirus proteases, including the PLRV protease. The non-functional *Rl_adg_* paralog *NLR_153_1* was used as a negative control. *Agrobacterium* (Agl1 strain) was infiltrated at OD_600_ = 0.3. Images were taken at 3 days post-infiltration. **(b)** Transient co-expression of *Rl_adg_* with infectious clones of PLRV or TuYV in *N. tabacum* also triggers HR. *Agrobacterium* (Agl1 strain) was infiltrated at OD_600_ = 0.3. The non-functional *Rl_adg_* paralog *NLR_153_1* was used as a negative control. Photographs were taken 4 days post-infiltration. **(c)** Overlaid structural predictions of the PLRV, TuYV and BMYV proteases show that despite amino acid divergence, these proteases share a conserved fold. **(d)** In *N. tabacum* transient assays, Rl_adg_ does not elicit HR when co-expressed with the proteases from CVEV, GEV-1, or PEMV-1. In contrast, co-expression with the PLRV protease produces clear cell death. The non-functional *Rl_adg_* paralog *NLR_153_1* was used as a negative control. *Agrobacterium* (Agl1 strain) was infiltrated at OD_600_ = 0.3. Photographs were taken 3 days post-infiltration.

The diversity of the tested poleroviruses is far greater than the variation between sequenced PLRV strains. The 22 published PLRV strains have proteases with at least 97.4% amino acid identity relative to the PLRV protease which was used in the transient expression assays (Supplementary Fig. 7). In contrast, the nine polerovirus proteases that elicited a response via Rl_adg_ shared between 40.6 and 82.6% amino acid identity to the PLRV protease (Supplementary Fig. 6). This suggests that Rl_adg_ is likely to recognise all published PLRV strains and could remain durable in the field.

To determine the limits of recognition, we tested Rl_adg_ against three proteases from the *Enamovirus* genus. Enamoviruses share a similar genome structure to poleroviruses and produce a self-cleaving P1 protein, but these proteases exhibit much lower sequence similarity to the PLRV protease (between 20 and 25%) (Supplementary Fig. 8a). Using Alphafold, we predicted the structures of three proteases from citrus vein enation virus (CVEV), grapevine enamovirus-1 (GEV-1) and pea enation mosaic virus-1 (PEMV-1). The predicted structures of these proteases were similar to those of polerovirus proteases (Supplementary Fig. 8b, Supplementary Fig. 8c). However, no cell death was observed when these three enamovirus proteases were co-expressed with Rl_adg_ in *N. tabacum* leaves (Fig. 4d). This suggests that the sequence or fold in polerovirus proteases recognised by Rl_adg_ is not conserved outside the *Polerovirus* genus, and the tested enamovirus proteases are too divergent to be recognised by Rl_adg_.

### Rl_adg_ associates with and oligomerises upon detection of polerovirus serine protease domains

To test for direct association between Rl_adg_ and the recognized protease domains, we performed a co-immunoprecipitation (CoIP) assay with FLAG-tagged Rl_adg_ and Myc-tagged protease domains co-expressed in the *eds1* mutant *N. benthamiana* line in which HR is abolished. Immunoprecipitated Rl_adg_ was analysed via SDS-PAGE and Western blotting. Association between Rl_adg_ and the PLRV protease was observed, but no interaction was detected between Rl_adg_ and the N-terminal P1 negative control protein. Similarly, Rl_adg_ co-immunoprecipitated with the TuYV and BMYV proteases (Fig. 5a).

**Figure 5.**
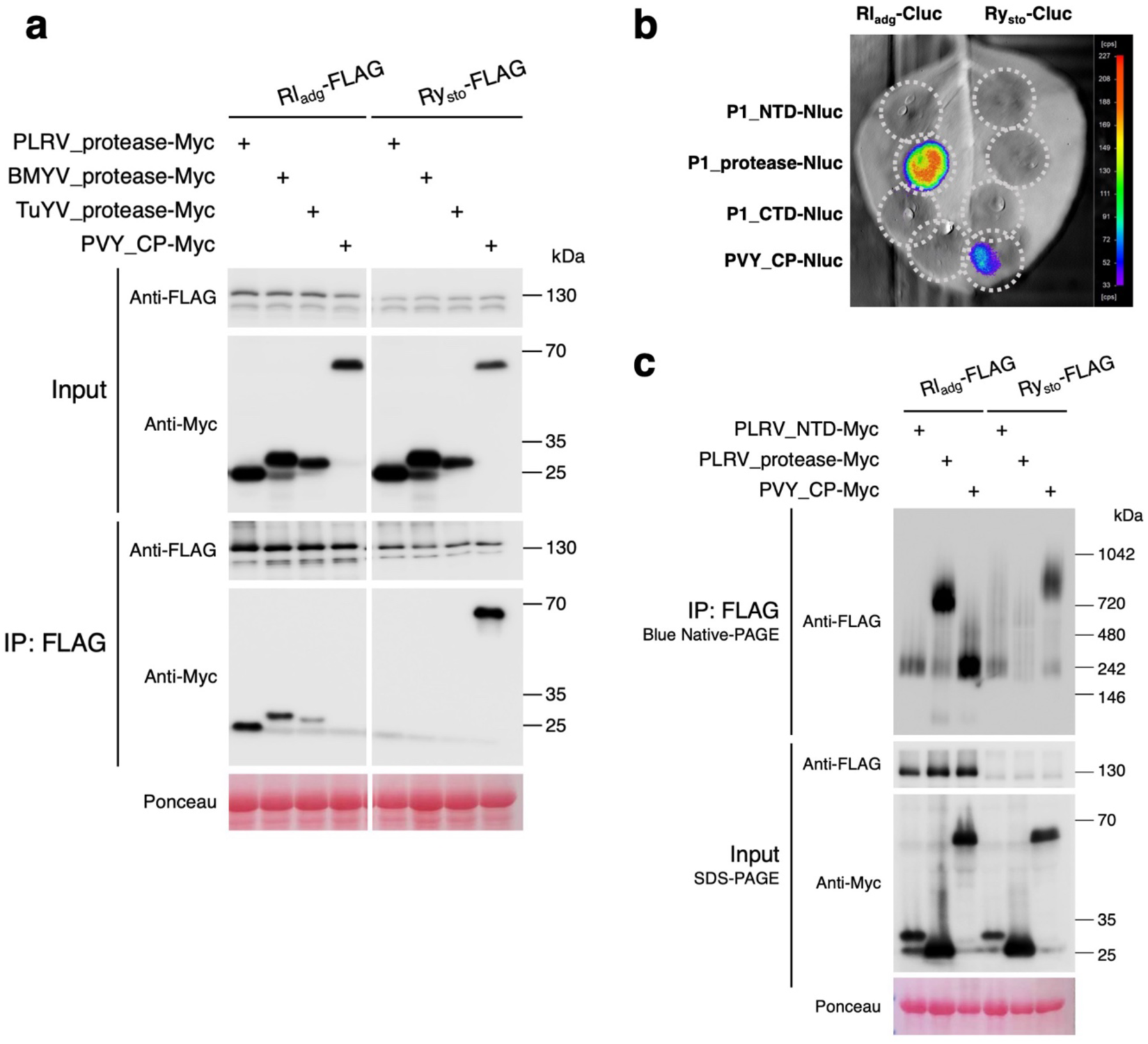
Rl_adg_ associates with, and oligomerizes upon detection of, polerovirus serine protease domains. **(a)** Proteases from PLRV, TuYV and BMYV co-precipitate with Rl_adg_. In contrast, the PVY coat protein does not co-precipitate with Rl_adg_. Ry_sto_ was used as a control, and PVY CP was successfully pulled down in a Ry_sto_-FLAG CoIP. Samples were prepared from agroinfiltrated leaves of the *eds1 N. benthamiana* line. *Agrobacterium* (Agl1 strain) was infiltrated at OD_600_ = 0.5, and samples were harvested 3 days post-infiltration. **(b)** A split-luciferase assay confirms *in planta* association between Rl_adg_ and the PLRV P1 protease. No luminescence signal was detected when *Rl_adg_* was co-expressed with the C-terminal or N-terminal domains of P1, or with the PVY CP. Co-expression of Ry_sto_ with the PVY CP served as a positive control. This assay was performed in the *eds1 N. benthamiana* line. *Agrobacterium* (Agl1 strain) was infiltrated at OD_600_ = 0.3. Leaves were imaged 3 days post-infiltration. **(c)** Rl_adg_ forms a higher-order oligomer upon co-expression with the P1 protease domain. FLAG-tagged Rl_adg_ or Ry_sto_ and Myc-tagged elicitors were co-expressed in the *N. benthamiana eds1* line. Leaf samples were harvested at 3 days post-infiltration, and protein complexes were separated on a blue native gel and visualized by immunoblotting to assess oligomerization state. *Agrobacterium* (Agl1 strain) was infiltrated at OD_600_ = 0.3. Samples were harvested and protein extracted 3 days post-infiltration.

To further investigate these interactions, we used a split luciferase assay. *Rl_adg_*, tagged with the C-terminal luciferase fragment, was co-expressed with the three P1 fragments tagged with the N-terminal luciferase fragment. Expression was performed in *eds1* mutant *N. benthamiana*. Luminescence was detected upon co-expression of *Rl_adg_* with the viral protease, but not with the other fragments of PLRV P1 (Fig. 5b). Combined, the split luciferase and CoIP demonstrate that Rl_adg_ recognises polerovirus proteases through direct interaction *in planta*, though the involvement of additional proteins is not ruled out This aligns with the HR observed for the HDS protease mutant, supporting a model of direct recognition rather than through an intermediate protein.

Next, non-denaturing native-PAGE was used to detect changes in the oligomeric state of Rl_adg_ upon protease detection. Co-expression of Rl_adg_ with the PLRV protease domain in *eds1 N. benthamiana* elicited formation of a slow-migrating Rl_adg_ oligomer. As expected, this was not observed when Rl_adg_ was expressed with the N-terminal P1 fragment or the PVY coat protein (Fig. 5c). Thus, like the TIR-NLRs ROQ1, RPP1, and Ry^sto^, Rl_adg_ transitions from a monomeric to an oligomeric form upon activation (Ma et al., 2020, Martin et al., 2020, Grech-Baran et al., 2025).

## Discussion

### *Rl_adg_* encodes a TIR-NLR which confers resistance to the phloem-limited PLRV

The polerovirus family encompasses diverse plant viruses and PLRV is an important constraint in seed potato production. Seed potato batches are rejected if PLRV levels exceed a threshold and control currently relies on chemical and cultural control of the aphid vector. To date, the only reported major *R*-gene against PLRV is *Rl_adg_,* found in the potato accession LOP-868. As *Rl_adg_* was previously mapped to an NLR-rich region on Chromosome 5 (Velasquez et al., 2007), we hypothesised that it encoded an NLR protein, likely belonging to either the *R1* or *Bs4* gene clusters.

Using resistance gene enrichment sequencing (RenSeq), combined with bulked segregant analysis, we identified NLR-encoding *Rl_adg_* candidates. Transient expression assays revealed *Rl_adg_* to be a *Bs4* ortholog encoding a TIR-NLR immune receptor. Given the rarity of genetic resistance to PLRV in cultivated potato, the identification of *Rl_adg_* provides a straightforward route to introgress resistance into susceptible varieties. Deploying this gene could reduce crop losses and decrease reliance on chemical control measures. However, many years of backcrossing and selection are required to introduce *Rl_adg_* into market-favoured potato varieties.

While NLR-encoding transcripts have been detected in phloem cells (Tang et al., 2023), Rl_adg_ is the first NLR found to confer resistance to a phloem-limited pathogen. This finding demonstrates that plants can mount an NLR-mediated immune response to infection within the vasculature. Notably, the P1 protease-triggered response in LOP-868 is detectable across the leaf, suggesting that Rl_adg_ function is not restricted to vascular tissue. *Rl_adg_* could thus confer resistance to PLRV during co-infection with other viruses such as PVY that enable PLRV to exit the vasculature (Barker, 1987, Savenkov and Valkonen, 2001).

### *Rl_adg_* confers resistance to many poleroviruses

Many NLRs recognize orthologous elicitors from multiple related pathogens, often by detecting a conserved structural feature. A well-characterised example is the PVY resistance protein Ry_sto_, which recognises the coat proteins of at least 10 potyviruses (Grech-Baran et al., 2022). Another is the *Capsicum annuum* CC-NLR Pvr4, which recognises the RNA-dependent RNA polymerases of multiple potyviruses (Kim et al., 2015, Kim et al., 2017). Beyond PLRV, several economically important poleroviruses threaten crop production, most notably TuYV in oilseed rape and BMYV, a component of the virus yellows complex in sugar beet. To determine whether Rl_adg_-mediated recognition extends to these pathogens, we used Alphafold to predict the structures of ten polerovirus proteases, including PLRV. All predicted models closely resembled each other and showed strong structural similarity to the published structure of the sesbania mosaic virus (SeMV) protease (Gayathri et al., 2006). When tested against a panel of these proteases, Rl_adg_ recognised all polerovirus variants despite their highly divergent amino acid sequences. However, no recognition was observed for three tested enamovirus proteases. These results suggest that, like Ry_sto_, Rl_adg_ detects multiple viruses via a conserved structural motif found across a virus family.

The breadth of Rl_adg_’s recognition capacity suggests it could be highly durable when deployed commercially in potato production. This durability is further supported by the low variability observed among PLRV strains and the protease’s essential role in polerovirus replication and maturation, making it unlikely that loss will enable resistance evasion. Our discovery that this recognition extends to other poleroviruses means that *Rl_adg_* could be transferred to other crop species - such as sugar beet – where the virus yellows complex poses a serious yield constraint.

### Rl_adg_ directly recognizes the serine protease structure

Rl_adg_ exhibits broad recognition of polerovirus serine proteases, despite the ten recognised proteases sharing little sequence similarity. We found that recognition is independent of protease catalytic activity because a catalytic mutant of the PLRV protease is also recognized. Co-immunoprecipitation and native-PAGE experiments indicate that Rl_adg_ activation occurs via direct association with the protease, triggering oligomerisation and initiating downstream immune signalling.

Although enamovirus proteases share a similar predicted fold to polerovirus proteases, co-expression with Rl_adg_ did not induce cell death. This lack of recognition may be due to subtle structural differences or specific changes within the region detected by Rl_adg_. High-resolution structural analysis - such as cryo-electron microscopy - could elucidate the interface between Rl_adg_ and the recognised proteases. This structural insight could enable engineering of Rl_adg_ to extend its recognition capacity. Structure-guided receptor engineering has proven successful for other NLRs, such as Pikp-1, Sr50, and MLA3 (Maidment et al., 2023, Tamborski et al., 2022, Gómez De La Cruz et al., 2026), where targeted amino acid substitutions in the binding interface expanded effector recognition, and also for the cell surface receptor FLS2 (Zhang et al., 2025, Li et al., 2025). Similar strategies could be used to enable detection of enamovirus proteases or to optimise the binding to specific polerovirus proteases, maximising recognition efficacy.

In summary, *Rl_adg_* is a *Bs4* ortholog encoding a TIR-NLR capable of broad-spectrum recognition of poleroviruses through direct detection of a conserved protease structure. Its wide recognition range and predicted durability make it an attractive candidate for engineering resistance across diverse plant species. With a structure-guided approach, Rl_adg_ specificity could be broadened beyond poleroviruses to target other plant pathogens that deploy serine proteases.

## Methods

### BSA-RenSeq and map-based cloning

Resistance gene enrichment sequencing (RenSeq) and bulked segregant sequencing was used to identify Rl_adg_ candidates. Genomic DNA from resistant and susceptible individuals were obtained from a previously characterised population (Velasquez et al., 2007). Enrichment for NLR-encoding genes was performed using the V4 RenSeq bait library (Witek et al., 2021). Libraries were sequenced using the paired 250 bp Illumina sequencing platform at The Earlham Institute.

Bulked segregant analysis (BSA) was performed following previously described methods (Jupe et al., 2013, Andolfo et al., 2014, Witek et al., 2016). Resistance-linked *R*-gene candidates were further filtered by their expression as determined using cDNA RenSeq and based on the presence of motifs identified using NLR-parser. The NLR genes on resistance-linked RenSeq contigs were predicted using NLR-parser (Steuernagel et al., 2015). To generate accurate gene models, cDNA RenSeq reads were mapped to reference sequences using the HISAT spliced aligner under default settings. Resulting BAM files were manually inspected to annotate NLR-encoding genes. Published sequences of the *R1* and *Bs4* were used to support the manual annotation.

### Constructs for transient expression and stable transformation

To facilitate testing of candidate genes, ORFs were initially cloned into binary vectors containing the CaMV 35S promoter and the *Agrobacterium tumefaciens* Ocs terminator. Candidates were cloned using either GoldenGate (into pICLS86922), or USER cloning (into pICSLUS0004OD) as previously described (Witek et al., 2016, Lin et al., 2023).

Additional constructs were generated to test *Rl_adg_* under its native regulatory elements. Constituent parts were cloned using the GoldenGate method using the vectors pICH41295 (promoter), pICH41308 (CDS), and pICH41276 (terminator). These parts were ligated into the binary vector pICH47742. To generate stably transformed *N. tabacum Native:Rl_adg_* lines, the binary vector pAGM31183 was used.

PLRV and TuYV infectious clones were used to assay resistance in *N. tabacum* transient assays. *Agrobacterium* infiltration was used to express these clones in leaf tissue. The infectious PLRV clone were obtained from Graham Cowan (James Hutton Institute) (Cowan et al., 2023). The TuYV infectious clone was supplied by Professor John Walsh (University of Warwick).

Transient expression assays for hypersensitive response (HR)

*Agrobacterium* suspensions were prepared to OD indicated in the figure legends. For HR assays in *N. tabacum,* fully expanded leaves were selected from plants aged between five and six weeks. For successful transient expression in *S. tuberosum*, plants were propagated in tissue culture, and rooted in 1% MS media for two weeks before being moved to compost. After another two weeks, leaves were infiltrated.

HR assays were performed by spot-infiltration of leaves using 1 ml needleless syringes. When multiple infiltrations were made on a single leaf, care was taken to avoid overlapping infiltration zones. Plants were left for 3-5 days to allow cell death to develop.

### Plant growth and transformation

*S. tuberosum* transformation was performed using previously described methods (Witek et al., 2016), the cultivar Maris Piper was used. For each transformation, a minimum of two independent lines is shown.

*N. benthamiana* was grown in a controlled environment room under conditions of 22 °C, 45-65% relative humidity, and a 16-hour light/ 8-hour dark cycle. Both wild type and *eds1* knockout *N. benthamiana* lines were used (Schultink et al., 2017).

### CoIP and blue native PAGE assays

Co-immunoprecipitation and native PAGE assays were conducted following previously described methods (Ahn et al., 2023). The C-terminal tags used in these experiments were; 3xFLAG (pICSL50007) for NLRs, and Myc (pICSL50010) for effectors. Protein was produced *in planta* by infiltrating whole *N. benthamiana* leaves with an *Agrobacterium* suspension, leaves were harvested 2 days post-infiltration. The NLRs were immunoprecipitated using a FLAG antibody, before visualisation using both FLAG and Myc antibodies.

### Split luciferase assay

To independently assess *in planta* association between NLRs and their cognate effectors, a split luciferase assay was performed as previously described (Lin et al., 2022).

C-terminal fragment of luciferase (pICSL50048) was fused to the C-terminus of Rl_adg_, while the PLRV P1 fragments were fused to an N-terminal luciferase fragment (pICSL50047). The binary vector pICSL86922 vector was used to express these fusions under the CaMV *35S* promoter and *Ocs* terminator. *Agrobacterium* strains carrying these constructs were spot-infiltrated into the *eds1 N. benthamiana* line. As a control for TIR-NLR interaction, the *S. stoloniferum* NLR Ry_sto_ and the potato virus Y coat protein were used.

Two days post-infiltration, leaves were harvested and infiltrated with 100 mM sodium citrate buffer containing 0.4 mM luciferin. Luminescence was visualised using the Nightowl II LB 983 In vivo imaging system and Winlight software (Berthold Technologies, Germany).

### Protein structure prediction, visualisation, and comparison

All structural predictions were run on the Alphafold 2 Colabfold webserver (Mirdita et al., 2022), confidence scores and pLDDT plots are presented in figures. Structural models were visualised in ChimeraX v1.6.1 (Meng et al., 2023). Structural similarity of protease structure models was assessed using DALI (Holm et al., 2023).

## Author contributions

R.H., K.W., and J.D.G.J. conceived and designed the project. R.H., H.Z., H-K.A., M.S., and K.W. performed the experiments. R.H., and K.W. performed the bioinformatic analyses. R.H. and J.D.G.J. wrote the manuscript with input from all authors. J.W., J.K., H.L-K., K.W., and J.D.G.J. contributed resources. All authors approved the manuscript.

## Acknowledgements

This research was supported by grants from the Gatsby Charitable Foundation to J.D.G.J. We thank the TSL transformation and tissue culture team led by Jodie Taylor, the SynBio team of Mark Youles and Liam Egan, media services, bioinformatics team, and the horticulturists (Sara Perkins, Justine Smith, Catherine Taylor, Timothy Wells, and Matt Castle) who supported this work. We also thank Ruth Norkor Annang for helpful comments on the manuscript.

## Supplementary Material

**Supplementary Figure 1.** *Rl_adg_* candidates are orthologs of the *R1* and *Bs4 R*-genes.

**Supplementary Figure 2.** *Rl_adg_* candidates from the *R1* gene family do not confer recognition of PLRV P1-P2.

**Supplementary Figure 3.** Native promoter-driven *Rl_adg_* confers PLRV recognition in *N. tabacum* transient expression assays.

**Supplementary Figure 4.** Rl_adg_ recognizes multiple polerovirus proteases despite their diverse amino acid sequences.

**Supplementary Figure 5.** Polerovirus proteases are predicted to share a conserved fold.

**Supplementary Figure 6.** Polerovirus proteases share conserved structural features despite low sequence identity.

**Supplementary Figure 7.** The P1 protease is highly conserved among published PLRV isolates.

**Supplementary Figure 8.** Enamovirus serine proteases have similar predicted structures to the PLRV protease despite low amino acid identity.

**Supplementary Table 1.** *Rl_adg_* is a *Bs4* orthologue

**Supplementary Table 2.** 22 PLRV sequences used in this work. **Supplementary Table 3.** Polerovirus isolate genomes used in this work. **Supplementary Table 4.** Three enamovirus sequences used in this work.

## References

Ahn, H.-K., Lin, X., Olave-Achury, A. C., Derevnina, L., Contreras, M. P., Kourelis, J., Wu, C.-H., Kamoun, S. & Jones, J. D. G. 2023. Effector-dependent activation and oligomerization of plant NRC class helper NLRs by sensor NLR immune receptors Rpi-amr3 and Rpi-amr1. The Embo Journal, 42, e111484.

Andika, I. B., Wei, S., Cao, C., Salaipeth, L., Kondo, H. & Sun, L. 2017. Phytopathogenic fungus hosts a plant virus: A naturally occurring cross-kingdom viral infection. Proceedings of the National Academy of Sciences, 114, 12267–12272.

Andolfo, G., Jupe, F., Witek, K., Etherington, G. J., Ercolano, M. R. & Jones, J. D. G. 2014. Defining the full tomato NB-LRR resistance gene repertoire using genomic and cDNA RenSeq. Bmc Plant Biology, 14.

Andret-Link, P., Schmitt-Keichinger, C., Demangeat, G., Komar, V. & Fuchs, M. 2004. The specific transmission of Grapevine fanleaf virus by its nematode vector Xiphinema index is solely determined by the viral coat protein. Virology, 320, 12–22.

Barker, H. 1987. Invasion of Non-phloem Tissue in Nicotiana clevelandii by Potato Leafroll Luteovirus Is Enhanced in Plants also Infected with Potato Y Potyvirus. Journal of General Virology, 68, 1223–1227.

Bendahmane, A., Kanyuka, K. & Baulcombe, D. C. 1999. The Rx gene from potato controls separate virus resistance and cell death responses. Plant Cell, 11, 781–92.

Bendahmane, A., Querci, M., Kanyuka, K. & Baulcombe, D. C. 2000. Agrobacterium transient expression system as a tool for the isolation of disease resistance genes: application to the Rx2 locus in potato. Plant J, 21, 73–81.

Bendix, C. & Lewis, J. D. 2018. The enemy within: phloem-limited pathogens. Mol Plant Pathol, 19, 238–254.

Carneiro, O. L. G., Ribeiro, S., Moreira, C. M., Guedes, M. L., Lyra, D. H. & Pinto, C. 2017. Introgression of the Rl(adg) allele of resistance to potato leafroll virus in Solanum tuberosum L. Crop Breeding and Applied Biotechnology, 17, 242–249.

Contreras, M. P., Lüdke, D., Pai, H., Toghani, A. & Kamoun, S. 2023. NLR receptors in plant immunity: making sense of the alphabet soup. EMBO reports, 24, e57495.

Cowan, G., Macfarlane, S. & Torrance, L. 2023. A new simple and effective method for PLRV infection to screen for virus resistance in potato. J Virol Methods, 315, 114691.

Esau, K. & Hoefert, L. L. 1972. Development of infection with beet western yellows virus in the sugarbeet. Virology, 48, 724–738.

Feehan, J. M., Castel, B., Bentham, A. R. & Jones, J. D. 2020. Plant NLRs get by with a little help from their friends. Curr Opin Plant Biol, 56, 99–108.

Gayathri, P., Satheshkumar, P. S., Prasad, K., Nair, S., Savithri, H. S. & Murthy, M. R. N. 2006. Crystal structure of the serine protease domain of Sesbania mosaic virus polyprotein and mutational analysis of residues forming the S1-binding pocket. Virology, 346, 440–451.

Gómez De La Cruz, D., Ingram, T., Zdrzałek, R., Taylor, J., Wawryk-Khamdavong, A., Bachowska, K., Banfield, M. J., Talbot, N. J. & Moscou, M. J. 2026. Molecular mimicry of a pathogen virulence target by a plant immune receptor. Science, 392, 1050–1055.

Grech-Baran, M., Witek, K., Ahn, H.-K., Lichocka, M., Vargas-Cortez, T., Barymow-Filoniuk, I., Witek, A. I., Hennig, J., Jones, J. D. G. & Poznański, J. T. 2025. Structural basis for heat tolerance in plant NLR immune receptors. bioRxiv, 2025.12.17.694812.

Grech-Baran, M., Witek, K., Jones, J. D. & Hennig, J. 2019. Coat protein of conserved structure from multiple potyviruses triggers Rysto-mediated immunity. Molecular Plant-Microbe Interactions, 32, 68–68.

Grech-Baran, M., Witek, K., Poznański, J. T., Grupa-URBAńska, A., Malinowski, T., Lichocka, M., Jones, J. D. G. & Hennig, J. 2022. The Rysto immune receptor recognises a broadly conserved feature of potyviral coat proteins. New Phytologist, 235, 1179–1195.

Greer, S. F., Hackenberg, D., Gegas, V., Mitrousia, G., Edwards, D., Batley, J., Teakle, G. R., Barker, G. C. & Walsh, J. A. 2021. Quantitative Trait Locus Mapping of Resistance to Turnip Yellows Virus in Brassica rapa and Brassica oleracea and Introgression of These Resistances by Resynthesis Into Allotetraploid Plants for Deployment in Brassica napus. Front Plant Sci, 12, 781385.

Hackenberg, D., Asare-Bediako, E., Baker, A., Walley, P., Jenner, C., Greer, S., Bramham, L., Batley, J., Edwards, D., Delourme, R., Barker, G., Teakle, G. & Walsh, J. 2020. Identification and QTL mapping of resistance to Turnip yellows virus (TuYV) in oilseed rape, Brassica napus. Theoretical and Applied Genetics, 133, 383–393.

Hameed, A., Iqbal, Z., Asad, S. & Mansoor, S. 2014. Detection of Multiple Potato Viruses in the Field Suggests Synergistic Interactions among Potato Viruses in Pakistan. Plant Pathol J, 30, 407–15.

Hemming, D., Bell, J., Collier, R., Dunbar, T., Dunstone, N., Everatt, M., Eyre, D., Kaye, N., Korycinska, A., Pickup, J. & Scaife, A. A. 2022. Likelihood of Extreme Early Flight of Myzus persicae (Hemiptera: Aphididae) Across the UK. Journal of Economic Entomology, 115, 1342–1349.

Holm, L., Laiho, A., TöRönen, P. & Salgado, M. 2023. Dali shines a light on remote homologs: One hundred discoveries. Protein Science, 32, e4519.

Jiang, Y., Zhang, C.-X., Chen, R. & He, S. Y. 2019. Challenging battles of plants with phloem-feeding insects and prokaryotic pathogens. Proceedings of the National Academy of Sciences, 116, 23390–23397.

Jingwei, L., Wang, B., Song, X.-M., Wang, R., Chen, L., Zhang, H., Hamborg, Z. & Wang, Q. 2013. Potato leafroll virus (PLRV) and Potato virus Y (PVY) influence vegetative growth, physiological metabolism, and microtuber production of in vitro-grown shoots of potato (Solanum tuberosum L.). Plant Cell, Tissue and Organ Culture (PCTOC), 114.

Juergens, M., Paetsch, C., Krämer, I., Zahn, M., Rabenstein, F., Schondelmaier, J., Schliephake, E., Snowdon, R., Friedt, W. & Ordon, F. 2010. Genetic analyses of the host-pathogen system Turnip yellows virus (TuYV)-rapeseed (Brassica napus L.) and development of molecular markers for TuYV-resistance. Theor Appl Genet, 120, 735–44.

Jupe, F., Pritchard, L., Etherington, G. J., Mackenzie, K., Cock, P. J., Wright, F., Sharma, S. K., Bolser, D., Bryan, G. J., Jones, J. D. & Hein, I. 2012. Identification and localisation of the Nb-Lrr gene family within the potato genome. BMC Genomics, 13, 75.

Jupe, F., Witek, K., Verweij, W., Sliwka, J., Pritchard, L., Etherington, G. J., Maclean, D., Cock, P. J., Leggett, R. M., Bryan, G. J., Cardle, L., Hein, I. & Jones, J. D. G. 2013. Resistance gene enrichment sequencing (RenSeq) enables reannotation of the NB-LRR gene family from sequenced plant genomes and rapid mapping of resistance loci in segregating populations. Plant Journal, 76, 530–544.

Kim, S.-B., Kang, W.-H., Huy, H. N., Yeom, S.-I., An, J.-T., Kim, S., Kang, M.-Y., Kim, H. J., Jo, Y. D., Ha, Y., Choi, D. & Kang, B.-C. 2017. Divergent evolution of multiple virus-resistance genes from a progenitor in Capsicum spp. New Phytologist, 213, 886–899.

Kim, S. B., Lee, H. Y., Seo, S., Lee, J. H. & Choi, D. 2015. Rna-dependent Rna polymerase (NIb) of the potyviruses is an avirulence factor for the broad-spectrum resistance gene Pvr4 in Capsicum annuum cv. CM334. PLos One, 10, e0119639.

Kondrák, M., Kopp, A., Uri, C., Sós-Hegedűs, A., Csákvári, E., Schiller, M., Barta, E., Cernák, I., Polgár, Z., Taller, J. & Bánfalvi, Z. 2020. Mapping and Dna sequence characterisation of the Rysto locus conferring extreme virus resistance to potato cultivar ’White Lady’. Plos one, 15, e0224534–e0224534.

Kuang, H., Wei, F., Marano, M. R., Wirtz, U., Wang, X., Liu, J., Shum, W. P., Zaborsky, J., Tallon, L. J., Rensink, W., Lobst, S., Zhang, P., Tornqvist, C. E., Tek, A., Bamberg, J., Helgeson, J., Fry, W., You, F., Luo, M. C., Jiang, J., Robin Buell, C. & Baker, B. 2005. The R1 resistance gene cluster contains three groups of independently evolving, type I R1 homologues and shows substantial structural variation among haplotypes of Solanum demissum. Plant J, 44, 37–51.

Lefeuvre, P., Martin, D. P., Elena, S. F., Shepherd, D. N., Roumagnac, P. & Varsani, A. 2019. Evolution and ecology of plant viruses. Nature Reviews Microbiology, 17, 632–644.

Li, T., Jarquin BOLAños, E., Stevens, D. M., Sha, H., Prigozhin, D. M. & Coaker, G. 2025. Unlocking expanded flagellin perception through rational receptor engineering. Nature Plants, 11, 1628–1641.

Lin, X., Jia, Y., Heal, R., Prokchorchik, M., Sindalovskaya, M., Olave-Achury, A., Makechemu, M., Fairhead, S., Noureen, A., Heo, J., Witek, K., Smoker, M., Taylor, J., Shrestha, R.-K., Lee, Y., Zhang, C., Park, S. J., Sohn, K. H., Huang, S. & Jones, J. D. G. 2023. Solanum americanum genome-assisted discovery of immune receptors that detect potato late blight pathogen effectors. Nature Genetics, 55, 1579–1588.

Lin, X., Olave-Achury, A., Heal, R., Pais, M., Witek, K., Ahn, H.-K., Zhao, H., Bhanvadia, S., Karki, H. S., Song, T., Wu, C.-H., Adachi, H., Kamoun, S., Vleeshouwers, V. G. A. A. & Jones, J. D. G. 2022. A potato late blight resistance gene protects against multiple Phytophthora species by recognizing a broadly conserved RXLR-WY effector. Molecular Plant, 15, 1457–1469.

Ma, S., Lapin, D., Liu, L., Sun, Y., Song, W., Zhang, X., Logemann, E., Yu, D., Wang, J., Jirschitzka, J., Han, Z., Schulze-Lefert, P., Parker, J. E. & Chai, J. 2020. Direct pathogen-induced assembly of an NLR immune receptor complex to form a holoenzyme. Science, 370, eabe3069.

Macleod, K., Greer, S. F., Bramham, L. E., Pimenta, R. J. G., Nellist, C. F., Hackenburg, D., Teakle, G. R., Barker, G. C. & Walsh, J. A. 2023. A review of sources of resistance to turnip yellows virus (TuYV) in Brassica species. Annals of Applied Biology, 183, 200–208.

Maidment, J. H. R., Shimizu, M., Bentham, A. R., Vera, S., Franceschetti, M., Longya, A., Stevenson, C. E. M., De La Concepcion, J. C., Białas, A., Kamoun, S., Terauchi, R. & Banfield, M. J. 2023. Effector target-guided engineering of an integrated domain expands the disease resistance profile of a rice NLR immune receptor. eLife, 12, e81123.

Martin, R., Qi, T., Zhang, H., Liu, F., King, M., Toth, C., Nogales, E. & Staskawicz, B. J. 2020. Structure of the activated ROQ1 resistosome directly recognizing the pathogen effector Xopq. Science, 370, eabd9993.

Meng, E. C., Goddard, T. D., Pettersen, E. F., Couch, G. S., Pearson, Z. J., Morris, J. H. & Ferrin, T. E. 2023. UCSF ChimeraX: Tools for structure building and analysis. Protein Science, 32, e4792.

Mihovilovich, E., Alarcon, L., Perez, A. L., Alvarado, J., Arellano, C. & Bonierbale, M. 2007. High levels of heritable resistance to Potato leafroll virus (PLRV) in Solanum tuberosum subsp andigena. Crop Science, 47, 1091–1103.

Mirdita, M., Schütze, K., Moriwaki, Y., Heo, L., Ovchinnikov, S. & Steinegger, M. 2022. ColabFold: making protein folding accessible to all. Nature Methods, 19, 679–682.

Nurkiyanova, K. M., Ryabov, E. V., Commandeur, U., Duncan, G. H., Canto, T., Gray, S. M., Mayo, M. A. & Taliansky, M. E. 2000. Tagging potato leafroll virus with the jellyfish green fluorescent protein gene. J Gen Virol, 81, 617–26.

Ryabov, E. V., Love, A. & Torrance, L. 2026. Potato leafroll virus: A re-emerging threat to sustainable potato production. Annals of Applied Biology, 189, e70140.

Sadowy, E., Juszczuk, M., David, C., Gronenborn, B. & Hulanicka, M. D. 2001. Mutational analysis of the proteinase function of Potato leafroll virus. J Gen Virol, 82, 1517–1527.

Savenkov, E. I. & Valkonen, J. P. T. 2001. Potyviral Helper-Component Proteinase Expressed in Transgenic Plants Enhances Titers of Potato Leaf Roll Virus but Does Not Alleviate Its Phloem Limitation. Virology, 283, 285–293.

Schultink, A., Qi, T., Lee, A., Steinbrenner, A. D. & Staskawicz, B. 2017. Roq1 mediates recognition of the Xanthomonas and Pseudomonas effector proteins XopQ and HopQ1. The Plant Journal, 92, 787–795.

Shepardson, S., Esau, K. & Mccrum, R. 1980. Ultrastructure of potato leaf phloem infected with potato leafroll virus. Virology, 105, 379–392.

Steuernagel, B., Jupe, F., Witek, K., Jones, J. D. G. & Wulff, B. B. H. 2015. NLR-parser: rapid annotation of plant Nlr complements. Bioinformatics, 31, 1665–1667.

Taliansky, M., Mayo, M. A. & Barker, H. 2003. Potato leafroll virus: a classic pathogen shows some new tricks. Mol Plant Pathol, 4, 81–9.

Tamada, T. & Kondo, H. 2013. Biological and genetic diversity of plasmodiophorid-transmitted viruses and their vectors. Journal of General Plant Pathology, 79, 307–320.

Tamborski, J., Seong, K., Liu, F., Staskawicz, B. & Krasileva, K., V. 2022. Engineering of Sr33 and Sr50 plant immune receptors to alter recognition specificity and autoactivity. bioRxiv, 2022.03.05.483131.

Tang, B., Feng, L., Hulin, M. T., Ding, P. & Ma, W. 2023. Cell-type-specific responses to fungal infection in plants revealed by single-cell transcriptomics. Cell Host Microbe, 31, 1732–1747.e5.

Velasquez, A. C., Mihovilovich, E. & Bonierbale, M. 2007. Genetic characterization and mapping of major gene resistance to potato leafroll virus in Solanumtuberosum ssp. andigena. Theor Appl Genet, 114, 1051–8.

Wang, K.-D., Dughbaj, M. A., Nguyen, T. T. V., Nguyen, T. Q. Y., Oza, S., Valdez, K., Anda, P., Waltz, J. & Sacco, M. A. 2023. Systematic mutagenesis of Polerovirus protein P0 reveals distinct and overlapping amino acid functions in Nicotiana glutinosa. Virology, 578, 24–34.

Wang, K.-D., Empleo, R., Nguyen, T. T. V., Moffett, P. & Sacco, M. A. 2015. Elicitation of hypersensitive responses in Nicotiana glutinosa by the suppressor of RNA silencing protein P0 from poleroviruses. Molecular Plant Pathology, 16, 435–448.

Witek, K., Jupe, F., Witek, A. I., Baker, D., Clark, M. D. & Jones, J. D. G. 2016. Accelerated cloning of a potato late blight-resistance gene using RenSeq and SMRT sequencing. Nature Biotechnology, 34, 656–660.

Witek, K., Lin, X., Karki, H. S., Jupe, F., Witek, A. I., Steuernagel, B., Stam, R., Van Oosterhout, C., Fairhead, S., Heal, R., Cocker, J. M., Bhanvadia, S., Barrett, W., Wu, C. H., Adachi, H., Song, T., Kamoun, S., Vleeshouwers, V., Tomlinson, L., Wulff, B. B. H. & Jones, J. D. G. 2021. A complex resistance locus in Solanum americanum recognizes a conserved Phytophthora effector. Nat Plants, 7, 198–208.

Zhang, S., Liu, S., Lai, H.-F., Bender, K. W., Kim, G., Caflisch, A. & Zipfel, C. 2025. Reverse engineering of the pattern recognition receptor FLS2 reveals key design principles of broader recognition spectra against evading flg22 epitopes. Nature Plants, 11, 1642–1657.

