## Supplementary figures and tables for "Broad-spectrum polerovirus resistance conferred by a potato TIR-NLR immune receptor"

Addresses:

[6.] 2Blades, Evanston, IL, 60201, USA

* Co-corresponding author

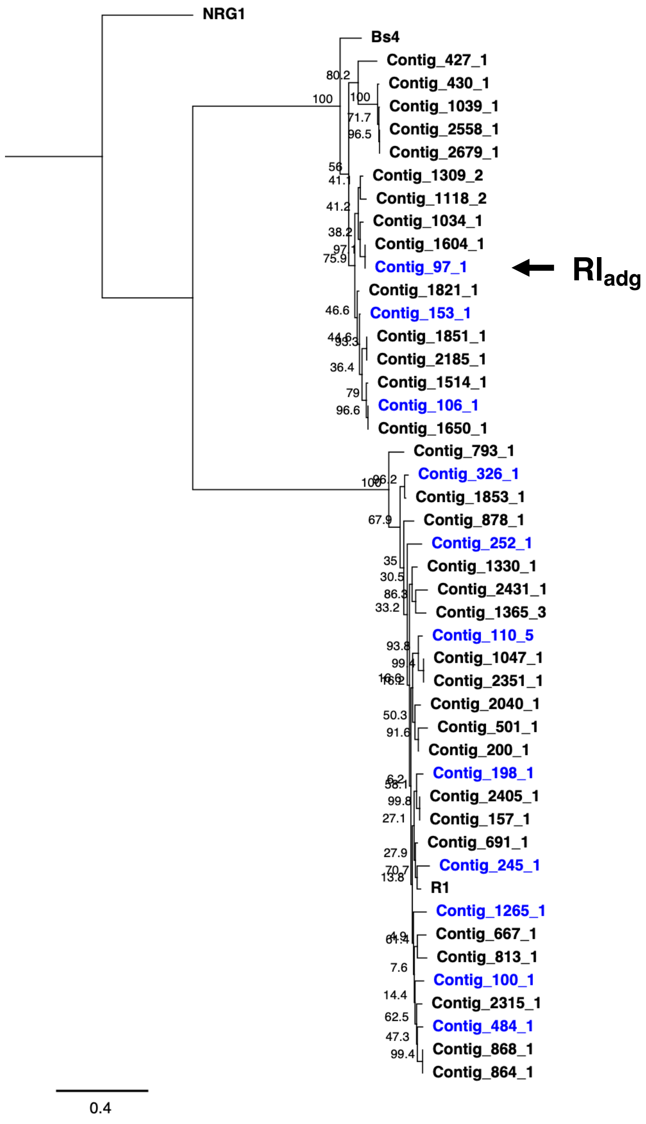


**Supplementary Figure 1. *Rl$_{adg}$* candidates are orthologs of the *R1* and *Bs4 R*-genes.** Bulked segregant analysis revealed that the *R1* and *Bs4* gene families co-segregate with PLRV resistance in LOP-868. In total, 27 R1-like and 17 Bs4-like NLRs were identified from PacBio RenSeq data. NLRs belonging to the resistant haplotype are highlighted in blue. Branch support values (%) are shown.

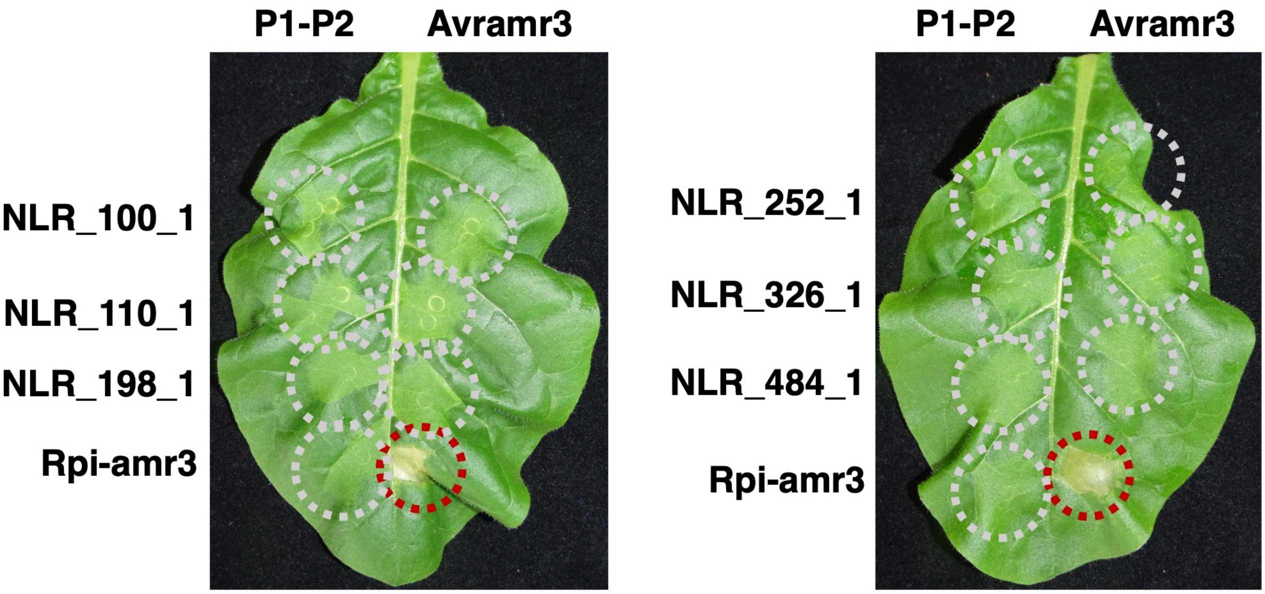


**Supplementary Figure 2. *$Rl_{adg}$* candidates from the *R1* gene family do not confer recognition of PLRV P1-P2.** Eight *R1* homologs are genetically linked to resistance in LOP-868; however, cDNA RenSeq data showed that only six are expressed. These six genes were cloned and co-expressed with the PLRV P1-P2 genomic region in *N. tabacum* leaves. No HR was observed in any infiltration. Avramr3 and Rpi-amr3 was used as controls. *Agrobacterium* (Agl1 strain) was infiltrated at $OD_{600}$ = 0.3, and images were taken 3 days post-infiltration.

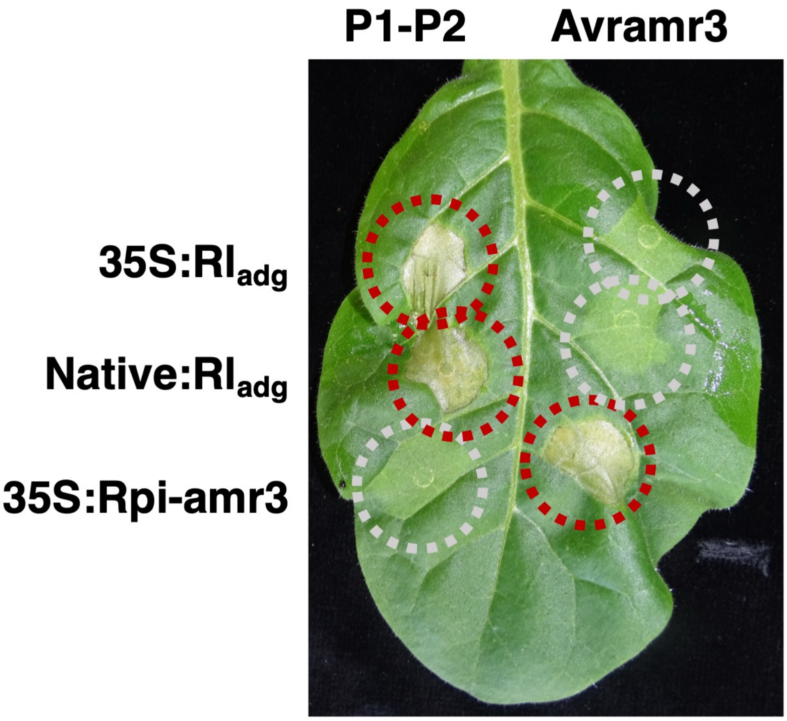


**Supplementary Figure 3. Native promoter-driven *Rl$_{adg}$* confers PLRV recognition in *N. tabacum* transient expression assays.** To demonstrate that *Rl$_{adg}$*-dependent HR in *N. tabacum* is not an artefact of overexpression, *Rl$_{adg}$* was cloned under its native regulatory elements. Co-expression of this construct with the PLRV P1-P2 region results in a clear HR. *Agrobacterium* (Agl1 strain) was infiltrated at $OD_{600} = 0.3$. Photographs were taken at 3 days post-infiltration.

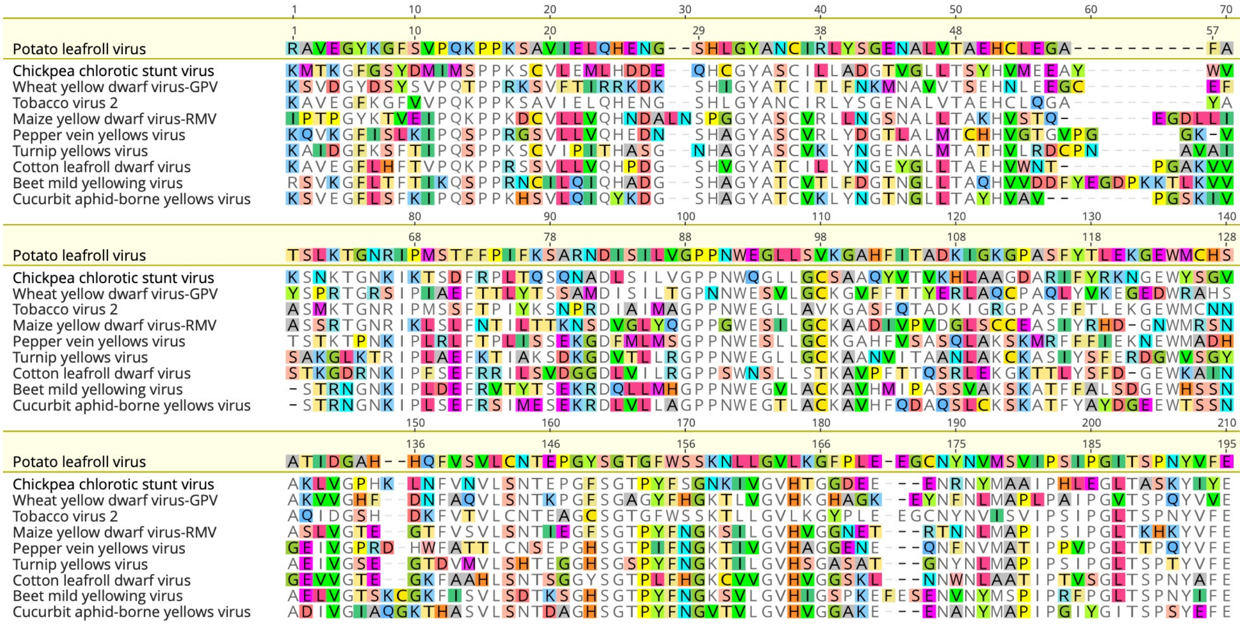


**Supplementary Figure 4. $Rl_{adg}$ recognizes multiple polerovirus proteases despite their diverse amino acid sequences.** Amino acid alignment of tested polerovirus proteases with the PLRV protease reveals limited sequence conservation. Polymorphic residues, relative to the PLRV reference, are highlighted. Despite this sequence divergence, all tested proteases are all recognized by $Rl_{adg}$.

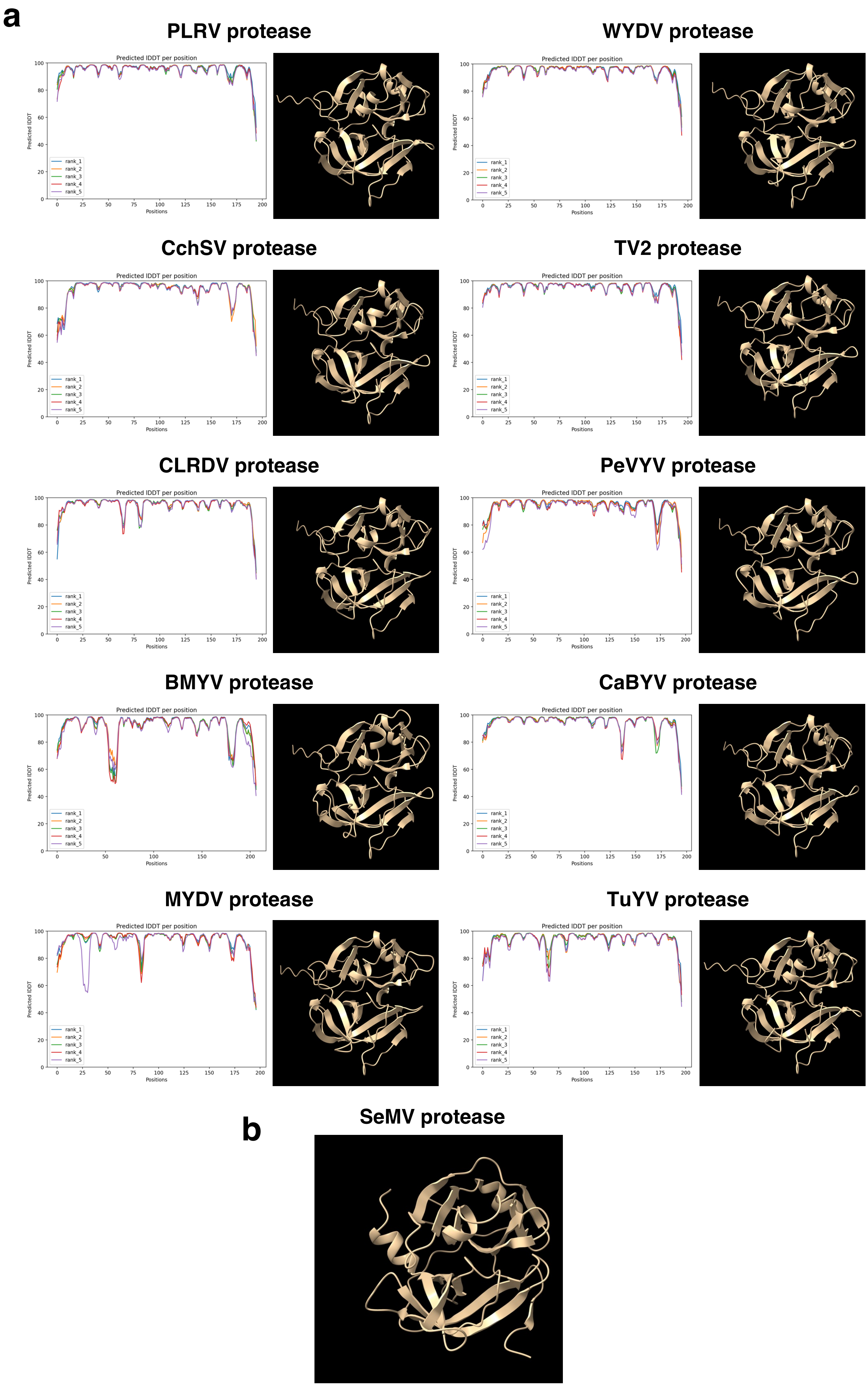
a
PLRV protease
WYDV protease
CchSV protease
TV2 protease
CLRDV protease
PeVYV protease
BMYV protease
CaBYV protease
MYDV protease
TuYV protease
Predicted lDDT per position
Predicted lDDT
Positions
rank_1
rank_2
rank_3
rank_4
rank_5
b
SeMV protease

**Supplementary Figure 5. Polerovirus proteases are predicted to share a conserved fold. (a)** AlphaFold was used to predict structures for all proteases tested against $Rl_{adg}$. Prediction confidence was evaluated using LDDT scores at each sequence position, which are plotted for each model. All proteases share a similar predicted structure, suggesting they likely adopt a fold similar to that of the PLRV protease. **(b)** The most closely related protease with an experimentally determined structure is the Sobemovirus SeMV protease, which adopts a chymotrypsin-like fold consistent with the AlphaFold predictions for polerovirus proteases.

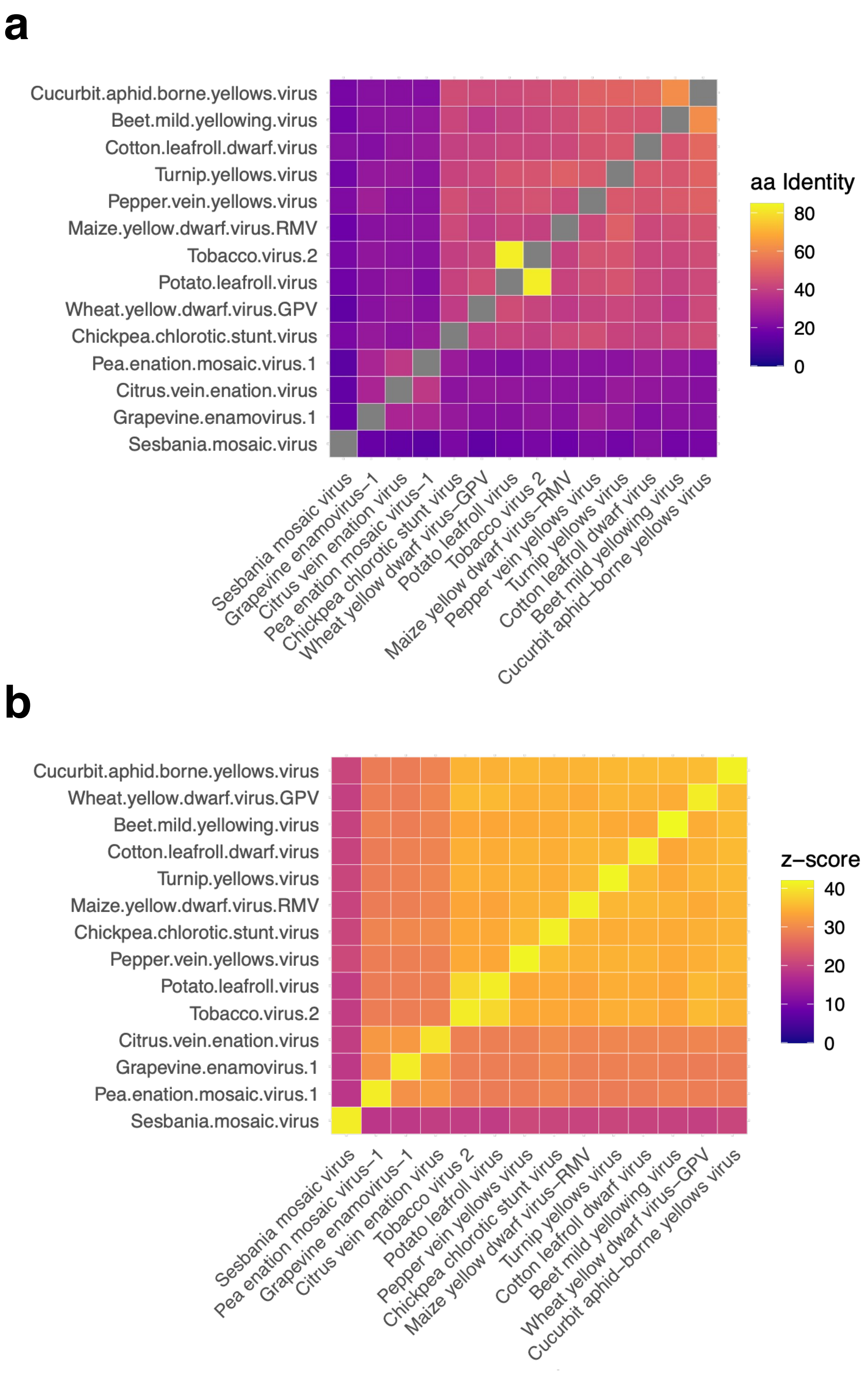


**Supplementary Figure 6. Polerovirus proteases share conserved structural features despite low sequence identity. (a)** Heatmap representing the pairwise amino acid identity for a selection of proteases from poleroviruses and enamoviruses. Polerovirus proteases tested against $Rl_{adg}$ share between 37.5% and 82.6% identity with the PLRV protease. **(b)** Despite low sequence identity, AlphaFold-predicted structures are highly similar. Pairwise structural comparisons were performed using DALI. z-scores from the DALI analysis are represented as a heatmap; scores above 20 indicate strong structural similarity.

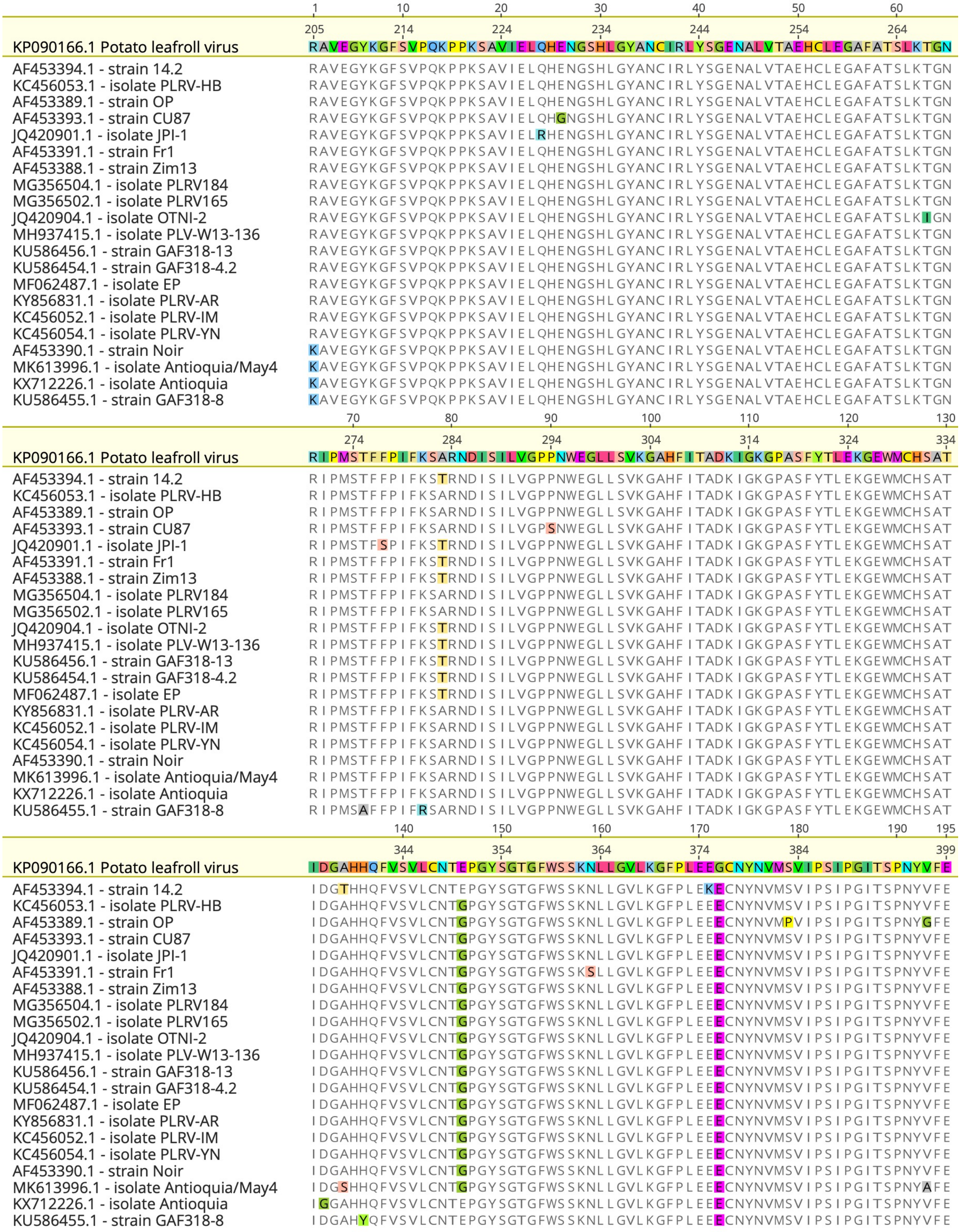


**Supplementary Figure 7. The P1 protease is highly conserved among published PLRV isolates.** All available PLRV genomes contain a P1 ORF and predicted protease. Amino acid identity between these alleles is high. The PLRV strain used for $Rl_{adg}$ identification and characterization (GenBank:KP090166.1) is included. Polymorphisms relative to the reference strain are highlighted in the alignment.

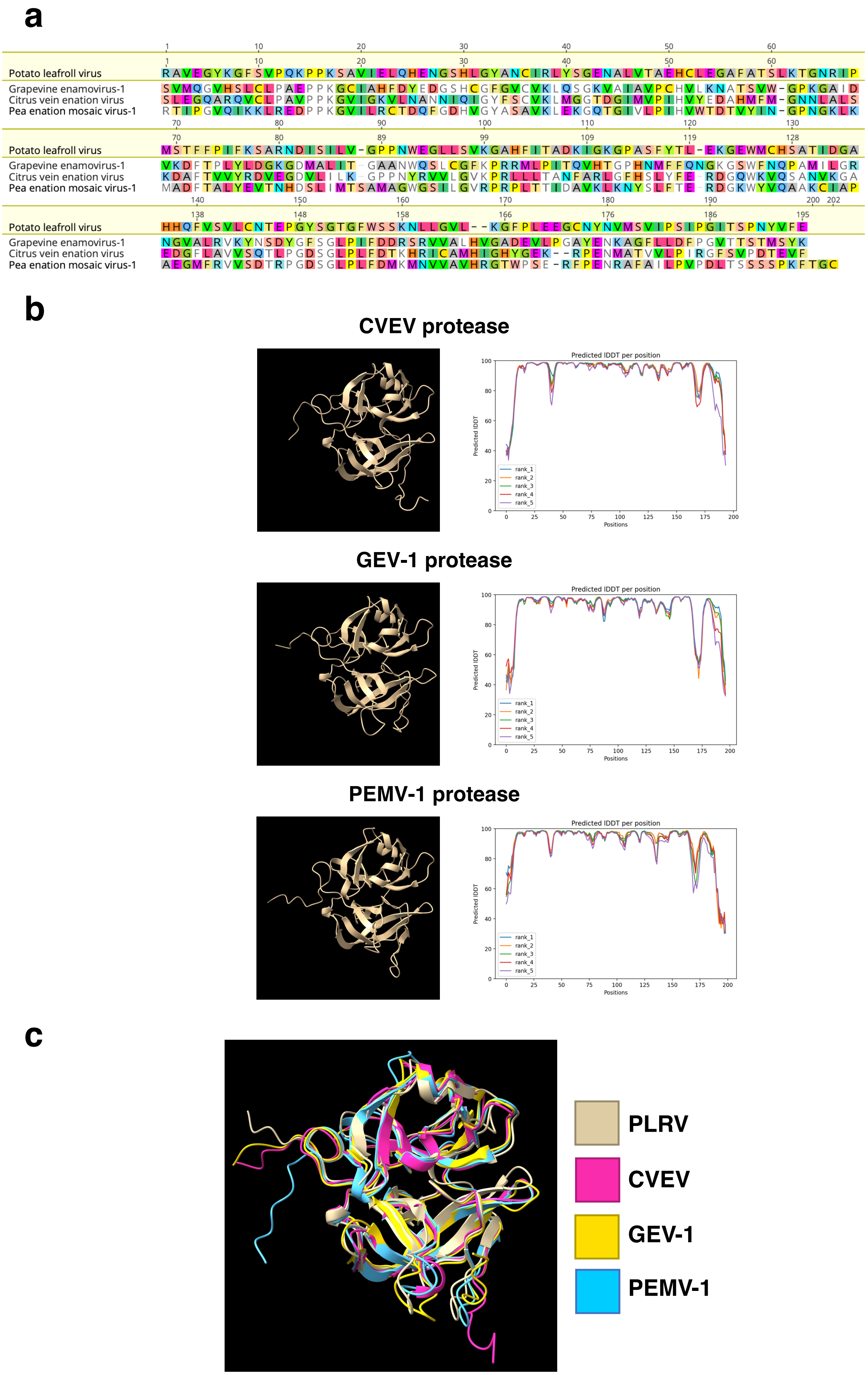
a
Potato leafroll virus
Grapevine enamovirus-1
Citrus vein enation virus
Pea enation mosaic virus-1
b
CVEV protease
GEV-1 protease
PEMV-1 protease
Predicted lDDT per position
Predicted lDDT
Positions
rank_1
rank_2
rank_3
rank_4
rank_5
c
PLRV
CVEV
GEV-1
PEMV-1

**Supplementary Figure 8. Enamovirus serine proteases have similar predicted structures to the PLRV protease despite low amino acid identity. (a)** Enamovirus proteases show low sequence similarity to the PLRV protease (20-25% amino acid identity). An alignment with the PLRV protease highlights polymorphisms in the enamovirus sequences. **(b)** AlphaFold was used to predict the structures of serine proteases from three enamoviruses: citrus vein enation virus (CVEV), grapevine enamovirus-1 (GEV-1), and pea enation mosaic virus-1 (PEMV-1). Prediction quality was assessed using LDDT scores across each sequence, which are shown for each model. **(c)** An overlay of the predicted structures demonstrates a highly similar fold to the PLRV protease. This structural similarity is supported by pairwise z-scores from DALI analysis (Supplementary figure 6b).

**Supplementary Table 1. *$Rl_{adg}$* is a *Bs4* orthologue.** *$Rl_{adg}$* maps to the *R1* and *Bs4 R*-gene clusters on Chromosome 5. Of the 11 NLRs identified on the resistant haplotypes of LOP-868, 8 were tested as $Rl_{adg}$ candidates. The ID, gene family assignment and expression in LOP-868 (determined using cDNA RenSeq of LOP-868) is shown. Cloned candidates were tested in *N. tabacum* by co-expression with the PLRV P1-P2 region, NLR_97_1 conferred gain of cell death in this assay.

| **NLR_ID** | **Gene family** | **Expression in LOP-868** | **HR with PLRV P1-P2** |
|---|---|---|---|
| NLR_100_1 | R1 | Yes | **No** |
| NLR_110_1 | R1 | Yes | **No** |
| NLR_198_1 | R1 | Yes | **No** |
| NLR_245_1 | R1 | **No** | - |
| NLR_252_1 | R1 | Yes | **No** |
| NLR_326_1 | R1 | Yes | **No** |
| NLR_484_1 | R1 | Yes | **No** |
| NLR_1265_1 | R1 | **No** | - |
| NLR_97_1 | Bs4 | Yes | **Yes** |
| NLR_106_1 | Bs4 | Yes | **Uncloned** |
| NLR_153_1 | Bs4 | Yes | **No** |

**Supplementary Table 2. 22 PLRV sequences used in this work.** The names and accession numbers for each of the 22 PLRV genomic sequences used in this work. The table also contains the URL to access each of these sequences.

| Strain ID | GenBank accession number | Link |
|---|---|---|
| Potato leafroll virus strain Zim13 | AF453388.1 | https://www.ncbi.nlm.nih.gov/nuccore/18656700 |
| Potato leafroll virus strain OP | AF453389.1 | https://www.ncbi.nlm.nih.gov/nuccore/AF453389.1 |
| Potato leafroll virus strain Noir | AF453390.1 | https://www.ncbi.nlm.nih.gov/nuccore/AF453390.1 |
| Potato leafroll virus strain Fr1 | AF453391.1 | https://www.ncbi.nlm.nih.gov/nuccore/AF453391.1 |
| Potato leafroll virus strain CU87 | AF453393.1 | https://www.ncbi.nlm.nih.gov/nuccore/AF453393.1 |
| Potato leafroll virus strain 14.2 | AF453394.1 | https://www.ncbi.nlm.nih.gov/nuccore/AF453394.1 |
| Potato leafroll virus isolate JPI-1 | JQ420901.1 | https://www.ncbi.nlm.nih.gov/nuccore/JQ420901.1 |
| Potato leafroll virus isolate OTNI-2 | JQ420904.1 | https://www.ncbi.nlm.nih.gov/nuccore/JQ420904.1 |
| Potato leafroll virus isolate PLRV-IM | KC456052.1 | https://www.ncbi.nlm.nih.gov/nuccore/KC456052.1 |
| Potato leafroll virus isolate PLRV-HB | KC456053.1 | https://www.ncbi.nlm.nih.gov/nuccore/KC456053.1 |
| Potato leafroll virus isolate PLRV-YN | KC456054.1 | https://www.ncbi.nlm.nih.gov/nuccore/KC456054.1 |
| Potato leafroll virus | KP090166.1 | https://www.ncbi.nlm.nih.gov/nuccore/KP090166.1 |
| Potato leafroll virus strain GAF318-4.2 | KU586454.1 | https://www.ncbi.nlm.nih.gov/nuccore/KU586454.1 |
| Potato leafroll virus strain GAF318-8 | KU586455.1 | https://www.ncbi.nlm.nih.gov/nuccore/KU586455.1 |
| Potato leafroll virus strain GAF318-13 | KU586456.1 | https://www.ncbi.nlm.nih.gov/nuccore/KU586456.1 |
| Potato leafroll virus isolate Antioquia | KX712226.1 | https://www.ncbi.nlm.nih.gov/nuccore/KX712226.1 |
| Potato leafroll virus isolate PLRV-AR | KY856831.1 | https://www.ncbi.nlm.nih.gov/nuccore/KY856831.1 |
| Potato leafroll virus isolate EP | MF062487.1 | https://www.ncbi.nlm.nih.gov/nuccore/MF062487.1 |
| Potato leafroll virus isolate PLRV165 | MG356502.1 | https://www.ncbi.nlm.nih.gov/nuccore/MG356502.1 |
| Potato leafroll virus isolate PLRV184 | MG356504.1 | https://www.ncbi.nlm.nih.gov/nuccore/MG356504.1 |
| Potato leafroll virus isolate PLV-W13-136 | MH937415.1 | https://www.ncbi.nlm.nih.gov/nuccore/MH937415.1 |
| Potato leafroll virus isolate Antioquia/May4 | MK613996.1 | https://www.ncbi.nlm.nih.gov/nuccore/MK613996.1 |

**Supplementary Table 3. Polerovirus isolate genomes used in this work.** The names, abbreviations and accession numbers for each of the 22 PLRV genomic sequences used in this work. The table also contains the URL to access each of these sequences. PLRV strains are not included but are shown in supplementary table 2.

| Strain ID | Abbreviated to | GenBank accession number | Accessible from |
|---|---|---|---|
| Beet mild yellowing virus isolate IPP | BMYV | DQ132996.1 | https://www.ncbi.nlm.nih.gov/nuccore/DQ132996 |
| Cucurbit aphid-borne yellows virus | CaBYV | EU000535.1 | https://www.ncbi.nlm.nih.gov/nuccore/EU000535 |
| Pepper vein yellows virus isolate HN | PeVYV | KP326573.1 | https://www.ncbi.nlm.nih.gov/nuccore/KP326573 |
| Turnip yellows virus isolate Landkreis Meissen_17 | TuYV | MN497810.1 | https://www.ncbi.nlm.nih.gov/nuccore/MN497810.1 |
| Chickpea chlorotic stunt virus | CchSV | NC_008249.1 | https://www.ncbi.nlm.nih.gov/nuccore/NC_008249 |
| Wheat yellow dwarf virus-GPV | WYDV | NC_012931.1 | https://www.ncbi.nlm.nih.gov/nuccore/NC_012931 |
| Cotton leafroll dwarf virus | CLRDV | NC_014545.1 | https://www.ncbi.nlm.nih.gov/nuccore/NC_014545 |
| Maize yellow dwarf virus-RMV | MYDV | NC_021484.1 | https://www.ncbi.nlm.nih.gov/nuccore/NC_021484 |
| Tobacco virus 2 | TV2 | NC_034265.1 | https://www.ncbi.nlm.nih.gov/nuccore/NC_034265 |

**Supplementary Table 4. Three enamovirus sequences used in this work.** The names, accession numbers and abbreviations for each of the enamovirus genomic sequences used in this work. The table also contains the URL to access each of these sequences.

| Strain ID | Abbreviated to | GenBank accession number | Accessible from |
|---|---|---|---|
| Grapevine enamovirus-1 isolate SE-BR | GEV-1 | NC_034836.1 | https://www.ncbi.nlm.nih.gov/nuccore/NC_034836.1 |
| Citrus vein enation virus, isolate VE-1 | CVEV | NC_021564.1 | https://www.ncbi.nlm.nih.gov/nuccore/NC_021564 |
| Pea enation mosaic virus-1 | PEMV | NC_003629.1 | https://www.ncbi.nlm.nih.gov/nuccore/NC_003629 |
